# Comparative transcriptome analysis provides insights into the resistance regulation mechanism and inhibitory effect of fungicide phenamacril in *Fusarium asiaticum*

**DOI:** 10.1101/2024.01.29.577693

**Authors:** Zhitian Zheng, Huaqi Liu, Xiao Luo, Runze Liu, Alexxander Joe, Haolin Li, Haiyan Sun, Lin Yanling, Yanzhong Li, Yunpeng Wang

## Abstract

*Fusarium asiaticum* is a destructive phytopathogenic fungus that causes Fusarium head blight of wheat (FHB), leading to serious yield and economic losses to cereal crops worldwide. Our previous studies indicated that target-site mutations (K216R/E, S217P/L, or E420K/G/D) of Type I myosin FaMyo5 conferred high resistance to phenamacril. Here, we first constructed a sensitive strain H1S and point mutation resistant strains HA, HC and H1R. Then we conducted comparative transcriptome analysis of these strains in *F. asiaticum* after 1 μg·mL^-1^ and 10 μg·mL^-1^ phenamacril treatment. Results indicated that 2135 genes were differentially expressed (DEGs) among the sensitive and resistant strains. Among them, the DEGs encoding ammonium transporter MEP1/MEP2, nitrate reductase, copper amine oxidase 1, 4-aminobutyrate aminotransferase, amino-acid permease inda1, succinate-semialdehyde dehydrogenase, 2, 3-dihydroxybenzoic acid decarboxylase, etc., were significantly up-regulated in all the phenamacril-resistant strains. Compared to the control group, a total of 1778 and 2097 DEGs were identified in these strains after 1 μg·mL^-1^ and 10 μg·mL^-1^ phenamacril treatment, respectively. These DEGs involved in 4-aminobutyrate aminotransferase, chitin synthase 1, multiprotein-bridging factor 1, transcriptional regulatory protein pro-1, amino-acid permease inda1, ATP-dependent RNA helicase DED, acetyl-coenzyme A synthetase, sarcoplasmic/endoplasmic reticulum calcium ATPase 2, etc., showed significantly down-regulated expression in phenamacril-sensitive strain but not in resistant strains after phenamacril treatment. In addition, cyanide hydratase, mating-type protein MAT-1, putative purine nucleoside permease, plasma membrane protein yro2, etc., showed significantly co-down-regulated expression in all the strains after phenamacril treatment. Taken together, This study provide deep insights into the resistance regulation mechanism and inhibitory effect of fungicide phenamacril and these new annotated proteins or enzymes are worth for the discovery of new fungicide targets.

**Author summary:** Fungicide phenamacril resistance occur in *F. asiaticum* and the resistance regulation mechanis are systematic and complex. Here, we conducted comparative transcriptome analysis of a sensitive strain H1S and point mutation resistant strains HA, HC and H1R in *F. asiaticum* after 1 μg·mL^-1^ and 10 μg·mL^-1^ phenamacril treatment. Among these annotated proteins or enzymes, amino-acid permease inda1, 1, 4-aminobutyrate aminotransferase, chitin synthase 1, multiprotein-bridging factor 1, ATP-dependent RNA helicase DED, acetyl-coenzyme A synthetase, sarcoplasmic/endoplasmic reticulum calcium ATPase 2, cyanide hydratase, mating-type protein MAT-1, putative purine nucleoside permease, plasma membrane protein yro2, etc., were involved in the resistance regulation mechanism and inhibitory effect of fungicide phenamacril. Our paper provides a reference basis for the study of drug resistance in other microorganisms. In addition, the relevant proteins or enzymes annotated in our study also have reference value for the discovery of new fungicide targets.

## Introduction

Fusarium head blight (FHB) or scab is a devastating disease of wheat, maize, barley, and other cereal crops worldwide [1]. Not only does this pathogen cause yield reductions, but also contaminate the grain with harmful trichothecene mycotoxins (e.g. deoxynivalenol, nivalenol, zearalenone) [2–4]. FHB is caused predominantly by fungi within the members of the *Fusarium graminearum* species complex (FGSC), which contains over 15 phylogenetic species (also known as lineages) [5–7]. Among these species, *F. asiaticum* and *F. graminearum* are the predominant causal agent of FHB in China [8,9]. Particularly, *F. asiaticum* is the major pathogenic population in the middle and lower reaches of the Yangtze River and the Huai River basin.

Because only few wheat cultivars are available with natural resistance to *Fusarium* spp. and complete genetic resistance to *F. asiaticum* has not been fully elucidated nor integrated into all commercial varieties [10], the use of fungicides during wheat flowering remain a very important tool for farmers in management of FHB in wheat in the last few decades [11,12]. Phenamacril (experimental code JS399-19; a.i. 2-cyano-3-amino-3-phenylancryic acetate), is a Fusarium-specific fungicide and shows excellent control of FHB and Rice bakanae disease in the field caused by *F. asiaticum* and *Fusarium fujikuroi*, respectively [13,14]. However, phenamacril-resistant mutants of *F. asiaticum* have been easily obtained by fungicide domestication and UV irradiation in the laboratory. Therefore, the Fungicide Resistance Action Committee (FRAC) classifies this fungicide as medium to high risk for resistance development (FRAC code list 2023, https://www.frac.info/docs/default-source/publications/frac-code-list/frac-code-list-2023---final.pdf).

We previously reported that phenamacril resistance in *Fusarium spp.* is caused by mutations in the Myosin5, encoded by FGSG_01410.1 [15]. In *F. asiaticum*, the mutation types of A135T, V151M, P204S, I434M, A577T, R580G/H or I581F led to low resistance to phenamacril [16]. The mutation types of S418R, I424R or A577G were responsible for moderate resistance and K216R/E, S217P/L or E420K/G/D conferred high resistance [16]. In *F. fujikuroi*, the point mutations K218T, S219P/L in Myosin-5 conferred high resistance to phenamacril in the field [14, 17]. In *Fusarium verticillioide*, single point mutations of S73L or E276K in the myosin-1 FvMyo1 were proven to be responsible for the high-level resistance to phenamacril [18]. In *Fusarium oxysporum*, FoMyo5 motor domain substitutions (V151A and S418T) cause natural low resistance to phenamacril [19]. In addition, Zhang et al., (2018) reported that point mutation S175L at myosin 5 conferred high resistance to phenamacril in *F. oxysporum* f. sp. *Melonis* [20]. On the other hand, previous studies revealed that phenamacril bound to FgMyoI, inhibited the ATPase activity or motor function of the wild-type FgMyo1 and had a serious effect on localization of FgMyo1 at the tips of germlings in *F. graminearum* [21–24].

In recent years, transcriptome analysis has become a powerful tool for understanding the development of organisms and the occurrence of diseases and exploring potential regulation pathways involved in pathogens response to fungicides stress [25–27]. Transcriptome sequencing is to study all mRNA transcribed by specific tissues or cells at a certain period or at a certain treatment [28]. High throughput transcriptome sequencing with its advantages of high sensitivity and wide application range has been widely used in the response mechanism of fungi to fungicides. For instance, the transcriptome analysis of *F. oxysporum* f. sp. *niveum* after treatment with 80 μg·mL^-1^ thymol, the gene transcription profiling of *F. graminearum* treated with an azole fungicide tebuconazole, the adaptation of *F. culmorum* to DMI fungicides and the comparative transcriptome analysis of *F. oxysporum* after 1 μg·mL^-1^ phenamacril treatment [24,25,29,30].

The objectives of this paper were to clarify the resistance regulation mechanism of phenamacril in *F. asiaticum* and determine the key genes or pathways where phenamacril could inhibit the growth of *F. asiaticum*. In this study, we first constructed a sensitive strain H1S and three point mutation resistant strains HA, HC and H1R (The types of point mutations are S217P, E420G and K216N, respectively). Then we conducted comparative transcriptome analysis of these phenamacril strains in *F. asiaticum* after 1 μg·mL^-1^ and 10 μg·mL^-1^ phenamacril treatment. These results could provide deeper insight into the resistance regulation mechanism and inhibitory effect of phenamacril in *F. asiaticum* and theoretical basis for the development of new fungicides to control FHB.

## Results

### Generation and identification of *F. asiaticum* FaMyo5 point mutantion strains by gene replacement strategy

In order to obtain phenamacril-sensitive strain and FaMyo5 point mutation phenamacril-resistant strains with consistent genetic background in *F. asiaticum*. Firstly, we introduced the *FaMyo5* gene fragments with point mutations S217P, E420G and K216N into the sensitive parental strain H1, respectively, by homologous double exchange, which we obtained the phenamacril-resistant strains HA, HC and H1R. Secondly, we inserted the trpC+HPH screening markers in front of the *FaMyo5* gene of the strain H1, which we obtained a phenamacril-sensitive strain H1S. The gene substitution process was shown in Fig 1A. Following purification by single-spore isolation, the gene replacement events were validated in the selected transformants by PCR and sequencing (Fig 1B).

**Fig 1.**
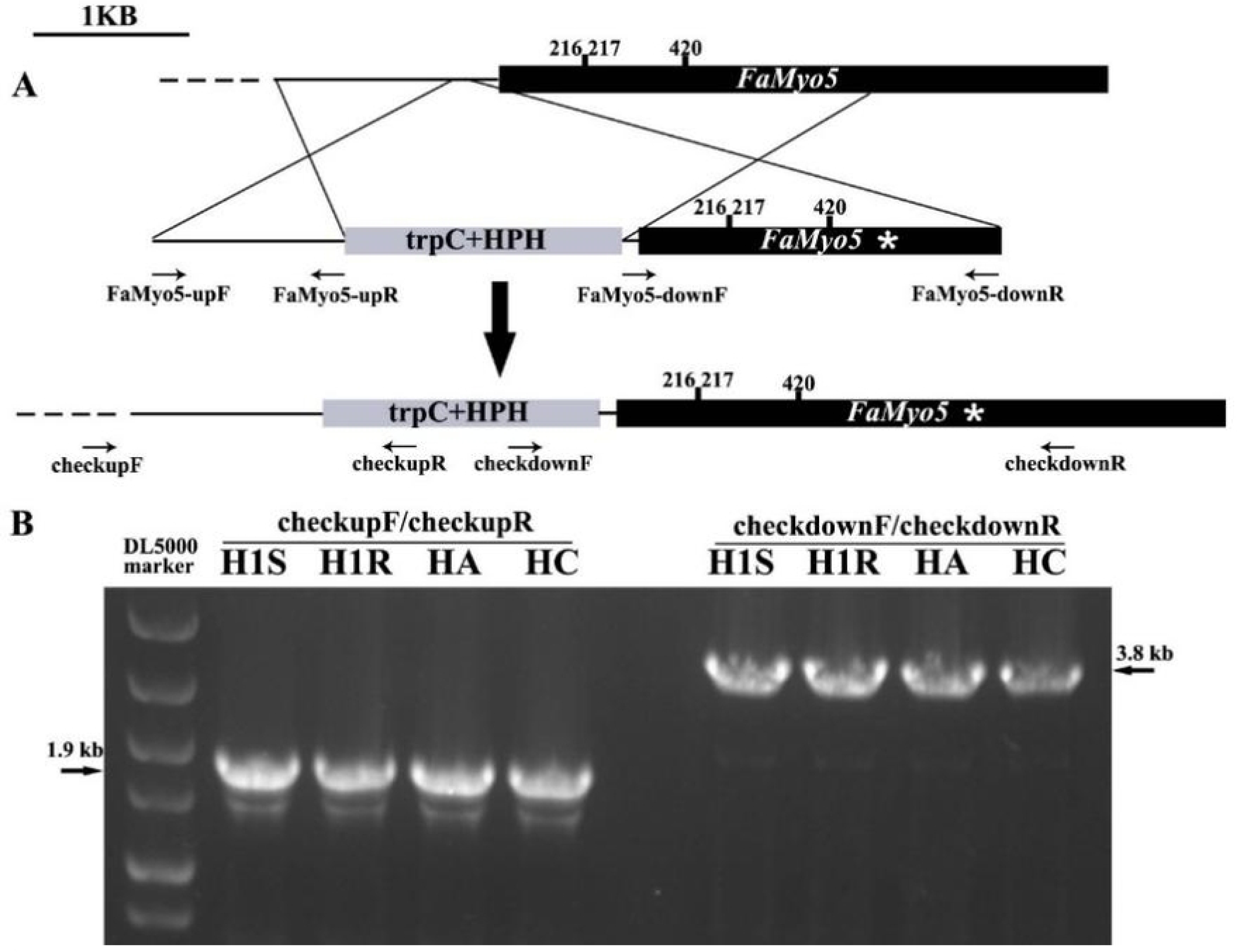
Generation and identification of phenamacril-sensitive strain and FaMyo5 point mutantion strains in *F. asiaticum*. (A) Schematic representation of gene replacement strategy. The upper part represents targeted gene replacement of *FaMyo5* was carried out using homologous double exchange. The black dashed lines represent the surrounding genomic region. The gene replacement cassette contains the hygromycin resistance gene (HPH) and the *FaMyo5* mutated zone. Primer binding sites are indicated by arrows (see S2 Table for the primer sequences). The asterisk represents the selected FaMyo5 mutated zone. Upon transformation, homologous recombination events take place (represented here by crosser over lines), resulting in replacement of the *FaMyo5* locus with HPH and mutated FaMyo5. (B) Polymerase chain reaction (PCR) strategy to screen replacement transformants. PCR performed with primer pairs checkupF/checkupR; a 1.9-kb amplified fragment indicates replacement integration at the left junction. PCR performed with primer pairs checkdownF/checkdownR; a 3.8-kb amplified fragment indicates replacement integration at the right junction.

In order to obtain the point mutation strains we designed, we aligned the FaMyo5 motor domains of the selected transformants using Bioedit 7.2 software. The sequences of *FaMyo5* motor domains were amplified from the cDNAs of *F. asiaticum* parental strain H1 and four mutant strains H1S, HA, HC, H1R following PCR amplification using primer pairs myosin5F/myosin5R (see Supporting information Table S2). The ORF of FaMyo5 motor domains encode a MYSc_Myo1 protein of 648 amino acids, which the coding region range from 52 to 699. Sequence analysis of the mutant strains confirmed that codons were altered as designed. For instance, in HA, codon 217 for serine S (TCA) was altered to a codon for proline P (CCA); In HC, codon 420 for glutamic acid E (GAA) was altered to a codon for glycine G (GGA); In H1R, codon 216 for lysine K (AAG) was altered to a codon for asparagine N (AAC); The FaMyo5 motor domain of strain H1S is completely consistent with that of the parental strain H1 (Fig 2A).

**Fig 2.**
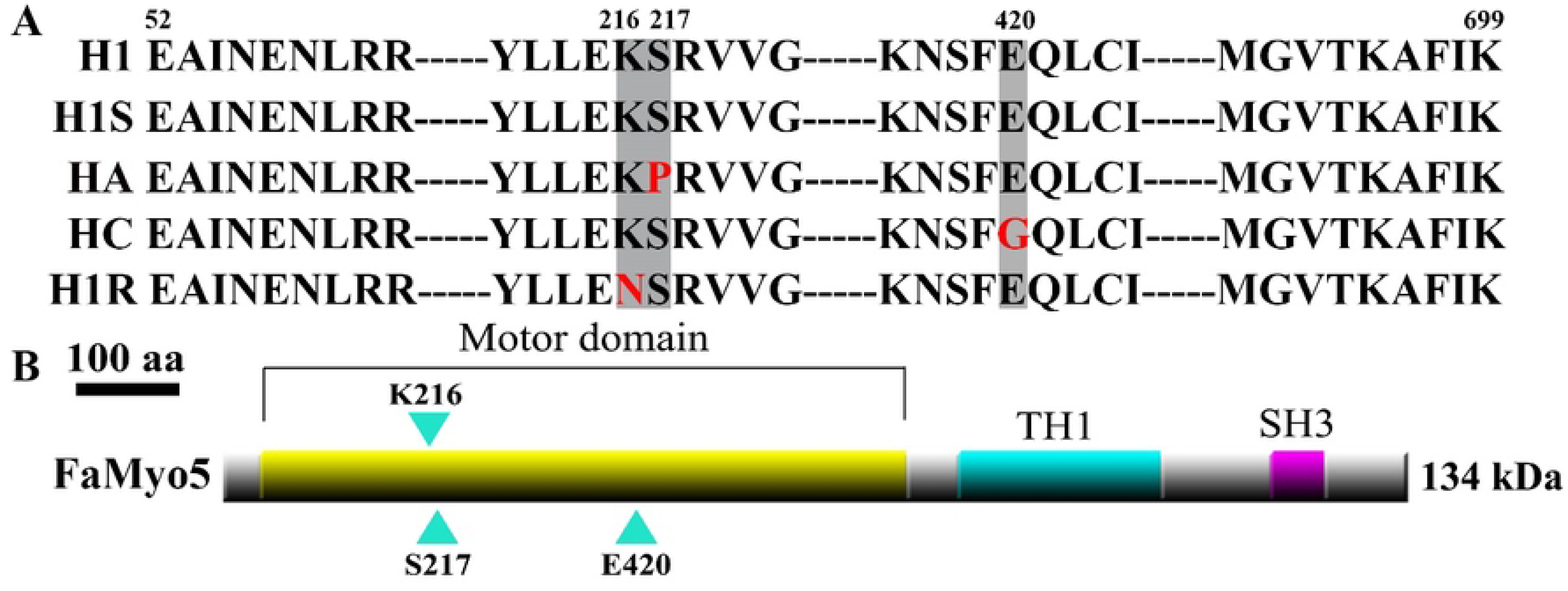
Analysis of point mutation types and domain of FaMyo5 in *F. asiaticum*. (A) Alignment of FaMyo5 motor domain from parental strain H1, phenamacril-sensitive strain H1S and point mutantion strains HA, HC, H1R. The vertical boxes indicate the amino acid changes at the codon 216, 217 and 420 that are responsible for phenamacril resistance. (B) Schematic representation of *F. asiaticum* FaMyo5. Sites of lysine, serine and glutamic acid mutantions are indicated with blue arrowheads. The conserved motor domain, myosin tail (TH1) and src homology domain 3 (SH3) are highlighted.

### The activity of phenamacril against *F. asiaticum* strains

We first compared the colony morphology and mycelial growth rate of *F. asiaticum* parental strain H1 and four mutant strains H1S, HA, HC, H1R. From the Fig 3 we can see, all of the strains exhibited white colonies. The mycelial growth rate of these *F. asiaticum* strains were not significantly changed and recorded in Table 1 in detail. Then we assessed the susceptibility of these *F. asiaticum* strains to phenamacril based on mycelial growth in fungicide-amended and fungicide-free media at 25 °C. Sensitivity test showed that phenamacril exhibits stronger inhibitory activity against sensitive strains H1 and H1S. The EC_50_ values of the two strains are 0.281 μg·mL^-1^ and 0.468 μg·mL^-1^, respectively (Table 1).

When treated with 10 μg·mL^-1^ phenamacril, they nearly could not grow on the PDA plates. However, all of the FaMyo5 point mutantion strains demonstrated high resistance levels to phenamacril, the EC_50_ values varied from 195.047 to 208.210 μg·mL^-1^ were recorded for HA, HC, and H1R (Table 1).

**Fig 3.**
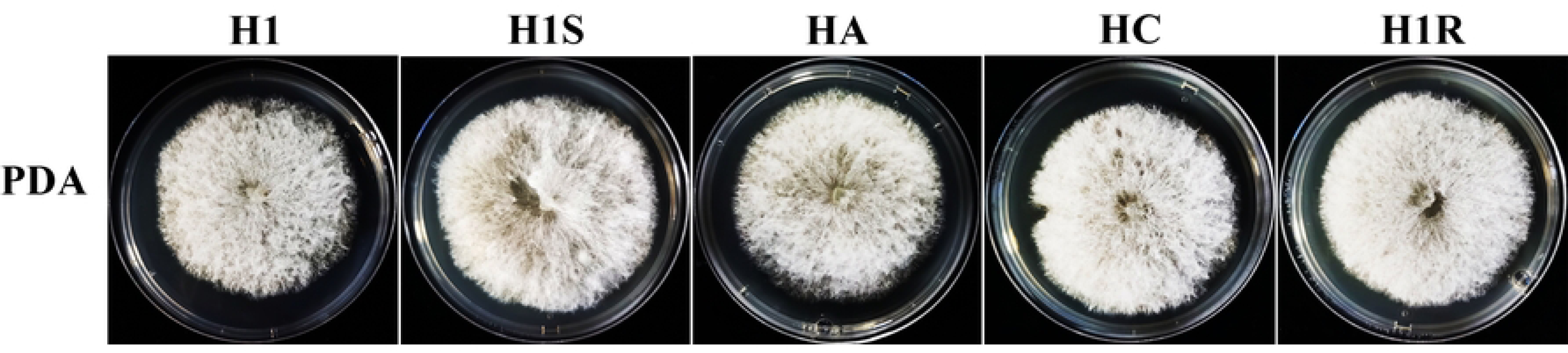
Colony morphology of *F. asiaticum* parental strain H1, phenamacril-sensitive strain H1S and point mutantion strains HA, HC, H1R. Strains were grown on solid media (PDA) for 3 days at 25 °C.

**Table 1.** Hyphal growth rate and sensitivity of the *F. asiaticum* strains to phenamacril.

| Strains | EC<sub>50</sub> ($\mu\text{g}\cdot\text{mL}^{-1}$)<sup>a</sup> | Growth rate on the media ($\text{mm}\cdot\text{day}^{-1}$)<sup>b</sup> |
| --- | --- | --- |
| H1 | 0.281 | 16.625±0.621a |
| H1S | 0.468 | 17.375±0.714a |
| HA | 208.210 | 17.500±0.816a |
| HC | 195.047 | 17.813±0.563a |
| H1R | 198.939 | 17.438±0.063a |

<sup>a</sup> EC<sub>50</sub> is the fungicide concentration resulting in 50% mycelial growth inhibition. Values are means of the experiments (differences among the experiments were not significant, P > 0.05, Fisher's LSD test).
<sup>b</sup> Growth rate was measured after incubation for 3 days. Mean and standard deviation were calculated from three replicates. The same letter indicated there was no significant difference.

### Transcriptome sequencing and data assembly in *F. asiaticum*

In order to elucidate the resistance regulation mechanism and inhibitory effect of phenamacril in *F. asiaticum*, we sequenced the transcriptomes of the phenamacril-resistant point mutantion strains HA, HC, H1R and one phenamacril-sensitive strain H1S following 12 h of 1 μg·mL^-1^ phenamacril or 10 μg·mL^-1^ phenamacril treatment and untreated strains using the Illumina Novaseq 6000 platform. 36 cDNA libraries were obtained, including H1S_CK, HA_CK, HC_CK, H1R_CK (untreated strains); H1S_1, HA_1, HC_1, H1R_1 (1 μg·mL^-1^ phenamacril treatment); H1S_10, HA_10, HC_10, H1R_10 (10 μg·mL^-1^ phenamacril treatment), and each treatment contained three biological repeats. A total of 28.44 to 35.97 million raw reads were generated. After removing adaptors and low-quality data, 21.86 to 34.38 million clean reads were obtained. Each library produced over 3G clean bases with a Q20 percentage over 98% (Table 2). When we compared these clean reads with the reference genome using HISAT2 software, our study found that over 99% of clean reads were uniquely mapped, while the proportion of multiple mapped reads was less than 1% (Table 2).

**Table 2.** Summary of statistical data for the transcriptome of *F. asiaticum*.

| Sample name | Raw reads | Clean reads | Clean bases | Q20 (%) | Q30 (%) | Mapped reads (%) | Uniquely mapped reads (%) | Multiple mapped reads (%) |
| --- | --- | --- | --- | --- | --- | --- | --- | --- |
| H1S_CK1 | 29360782 | 24574656 | 3442079437 | 98.89 | 95.74 | 90.47 | 99.64 | 0.36 |
| H1S_CK2 | 29241480 | 25860426 | 3623857224 | 98.89 | 95.72 | 90.80 | 99.58 | 0.42 |
| H1S_CK3 | 31688760 | 29221300 | 4099311236 | 98.87 | 95.68 | 90.93 | 99.49 | 0.51 |
| H1S_1_1 | 30611316 | 27673936 | 3878852546 | 98.89 | 95.77 | 90.24 | 99.56 | 0.44 |
| H1S_1_2 | 29150092 | 24524908 | 3435067764 | 98.92 | 95.82 | 90.21 | 99.53 | 0.47 |
| H1S_1_3 | 31676708 | 28263796 | 3961579174 | 98.91 | 95.79 | 90.90 | 99.52 | 0.48 |
| H1S_10_1 | 31223664 | 28083568 | 3928765399 | 98.97 | 96.02 | 89.80 | 99.57 | 0.43 |
| H1S_10_2 | 28970092 | 25161904 | 3525124928 | 98.91 | 95.8 | 89.71 | 99.52 | 0.48 |
| H1S_10_3 | 32042972 | 28534090 | 3997137960 | 98.9 | 95.77 | 89.59 | 99.52 | 0.48 |
| HA_CK1 | 29749344 | 26726674 | 3748707772 | 98.89 | 95.74 | 90.87 | 99.67 | 0.33 |
| HA_CK2 | 28970494 | 24874548 | 3484735535 | 98.96 | 95.98 | 90.52 | 99.51 | 0.49 |
| HA_CK3 | 29279812 | 25592552 | 3585507974 | 98.96 | 95.96 | 90.64 | 99.49 | 0.51 |
| HA_1_1 | 29923454 | 26198784 | 3673419928 | 98.91 | 95.79 | 90.76 | 99.54 | 0.46 |
| HA_1_2 | 28440464 | 21859752 | 3060148864 | 98.95 | 95.9 | 90.40 | 99.55 | 0.45 |
| HA_1_3 | 28955848 | 25566002 | 3583675523 | 98.93 | 95.86 | 90.88 | 99.52 | 0.48 |
| HA_10_1 | 35968074 | 34380366 | 4821052550 | 98.82 | 95.59 | 90.98 | 99.53 | 0.47 |
| HA_10_2 | 30268582 | 25859010 | 3623300524 | 98.91 | 95.79 | 90.48 | 99.48 | 0.52 |
| HA_10_3 | 31156754 | 25797074 | 3616490283 | 98.84 | 95.58 | 90.94 | 99.52 | 0.48 |
| HC_CK1 | 31434356 | 27082236 | 3797465208 | 98.88 | 95.69 | 90.37 | 99.63 | 0.37 |
| HC_CK2 | 32651444 | 29832572 | 4181403115 | 98.9 | 95.78 | 90.53 | 99.61 | 0.39 |
| HC_CK3 | 32574652 | 28376550 | 3973354503 | 98.9 | 95.76 | 90.09 | 99.61 | 0.39 |
| HC_1_1 | 32947828 | 30377880 | 4258166348 | 98.87 | 95.65 | 89.54 | 99.60 | 0.40 |
| HC_1_2 | 30890224 | 27868994 | 3906361880 | 98.89 | 95.73 | 89.94 | 99.58 | 0.42 |
| HC_1_3 | 32092812 | 29446944 | 4128766160 | 98.86 | 95.67 | 90.36 | 99.54 | 0.46 |
| HC_10_1 | 30320164 | 25602918 | 3586067287 | 98.87 | 95.67 | 89.57 | 99.59 | 0.41 |
| HC_10_2 | 28837894 | 23295018 | 3262268751 | 98.9 | 95.79 | 89.53 | 99.58 | 0.42 |
| HC_10_3 | 31823520 | 28930674 | 4058090144 | 98.84 | 95.59 | 90.73 | 99.62 | 0.38 |
| H1R_CK1 | 32978448 | 29558386 | 4142939796 | 98.91 | 95.81 | 90.85 | 99.62 | 0.38 |
| H1R_CK2 | 32586690 | 28730492 | 4026140831 | 98.89 | 95.73 | 90.61 | 99.65 | 0.35 |
| H1R_CK3 | 34020904 | 31144134 | 4366603481 | 98.84 | 95.59 | 90.29 | 99.55 | 0.45 |
| H1R_1_1 | 32233164 | 29445412 | 4130300119 | 98.86 | 95.65 | 90.20 | 99.66 | 0.34 |
| H1R_1_2 | 31443378 | 27923954 | 3916012148 | 98.86 | 95.67 | 90.00 | 99.51 | 0.49 |
| H1R_1_3 | 33037300 | 30482410 | 4274380891 | 98.84 | 95.57 | 90.24 | 99.56 | 0.44 |
| H1R_10_1 | 31281268 | 27397384 | 3840977496 | 98.85 | 95.61 | 89.67 | 99.60 | 0.40 |
| H1R_10_2 | 31802020 | 27288362 | 3826748502 | 98.88 | 95.69 | 89.56 | 99.64 | 0.36 |
| H1R_10_3 | 33128504 | 29921920 | 4196986035 | 98.82 | 95.53 | 89.92 | 99.62 | 0.38 |

A total of 13,313 unigenes were obtained from 36 high-quality transcriptomes, and the abundance of all unigenes was standardized and calculated by FPKMs (FPKM, expected number of Fragments Per Kilobase of transcript sequence per Millions base pairs sequenced) using uniquely mapped reads. In addition, functional annotation of all the unigenes were conducted and a total of 13,313 unigenes were annotated by searching the NCBI, Uniprot, GO, and KEGG databases, respectively. Therefore, this high-quality transcriptome sequencing represents a valuable resource for further research on *F. asiaticum* strains.

### Analysis of differential expression genes (DEGs) after phenamacril treatment

One of the primary goals of the transcriptome study was to identify variations between phenamacril-sensitive and -resistant strains after phenamacril treatment. The results indicate that these variations ranged from 114 to 817 DEGs, based on the FPKM value (Fig 4). All DEGs conformed to the absolute value of |log_2_FC| >1 and FDR < 0.05. Compared with the phenamacril-sensitive strain H1S, there were 669, 499, and 488 DEGs significantly up-regulated expression and 379, 450, and 817 DEGs significantly down-regulated expression in phenamacril-resistant strains HA, HC, and H1R, respectively. When treated with 1 μg·mL^-1^ phenamacril, there were 308 DEGs were significantly up-regulated and 465 DEGs were significantly down-regulated in phenamacril-sensitive strain H1S, while 200, 224, and 234 DEGs were significantly up-regulated and 381, 337, and 260 DEGs were significantly down-regulated in phenamacril-resistant strains HA, HC, and H1R, respectively. When treated with 10 μg·mL^-1^ phenamacril, 543 DEGs were significantly up-regulated and 473 DEGs were significantly down-regulated in phenamacril-sensitive strain H1S, while 114, 168, and 166 DEGs were significantly up-regulated and 418, 327, and 497 DEGs were significantly down-regulated in phenamacril-resistant strains HA, HC, and H1R, respectively (Fig 4).

**Fig 4.**
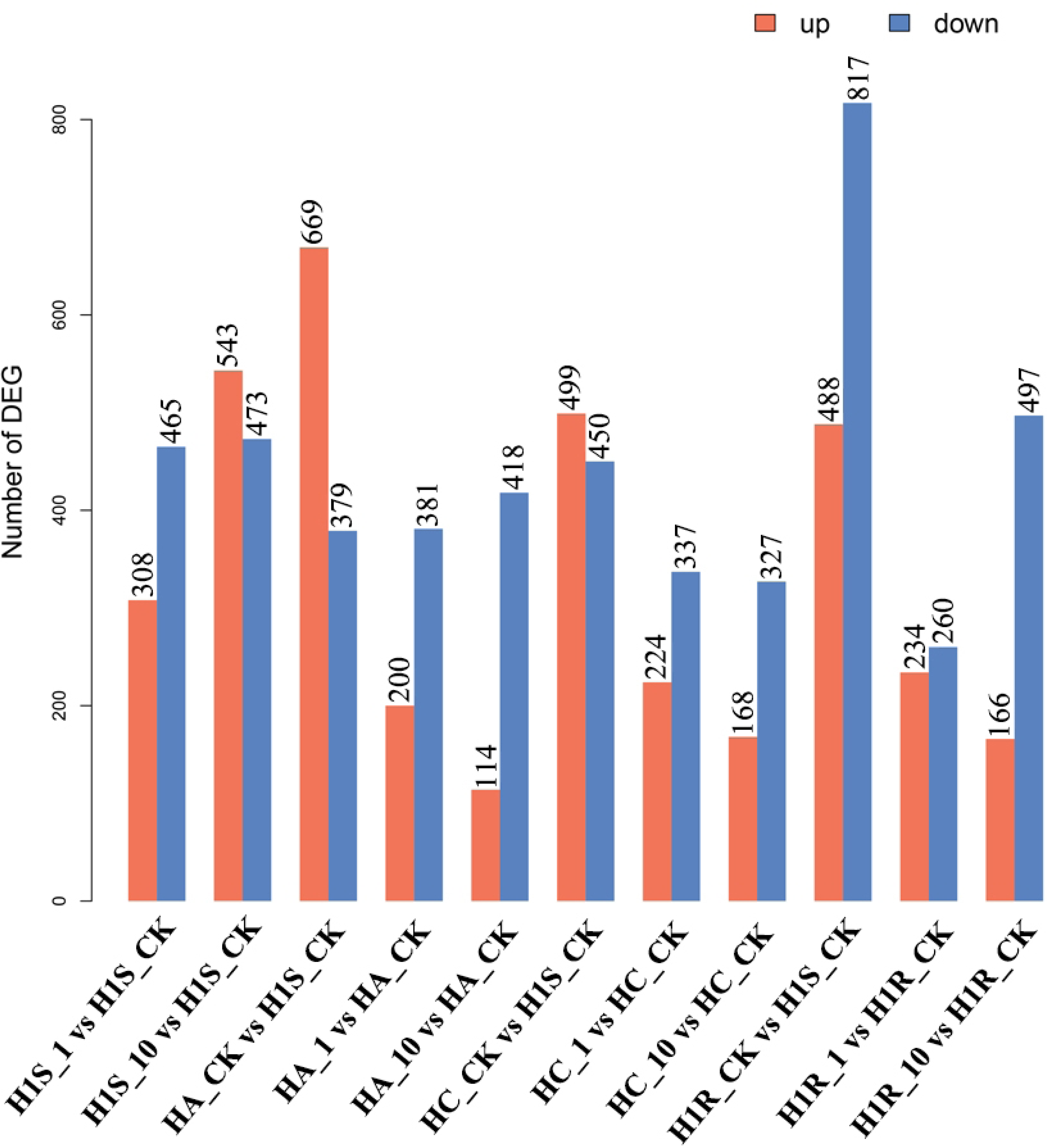
Analysis of DEGs in *F. asiaticum*. The red column indicates up-regulated expression, while the blue column indicates down-regulated expression.The abscissa represents the experimental group and the ordinate represents the number of DEGs.

The relationships among different DEG groups from Fig. 5 were displayed as Venn diagrams, and the results indicated that 276 DEGs were identified in three phenamacril-resistant strains when compared with the phenamacril-sensitive strain H1S (Fig 5C). When treated with 1 μg·mL^-1^ phenamacril, 18 DEGs were identified in both phenamacril-resistant and -sensitive strains; 16 DEGs were identified in all the resistant strains and 462 DEGs were only identified in the sensitive strain H1S (Fig 5A). When treated with 10 μg·mL^-1^ phenamacril, 5 DEGs were identified in both phenamacril-resistant and -sensitive strains; 11 DEGs were identified in all the resistant strains and 683 DEGs were identified only in the sensitive strain H1S (Fig 5B). By further studying the function of these DEGs, we might find the mechanism of regulating resistance and response stress to phenamacril in *F. asiaticum*.

**Fig. 5.**
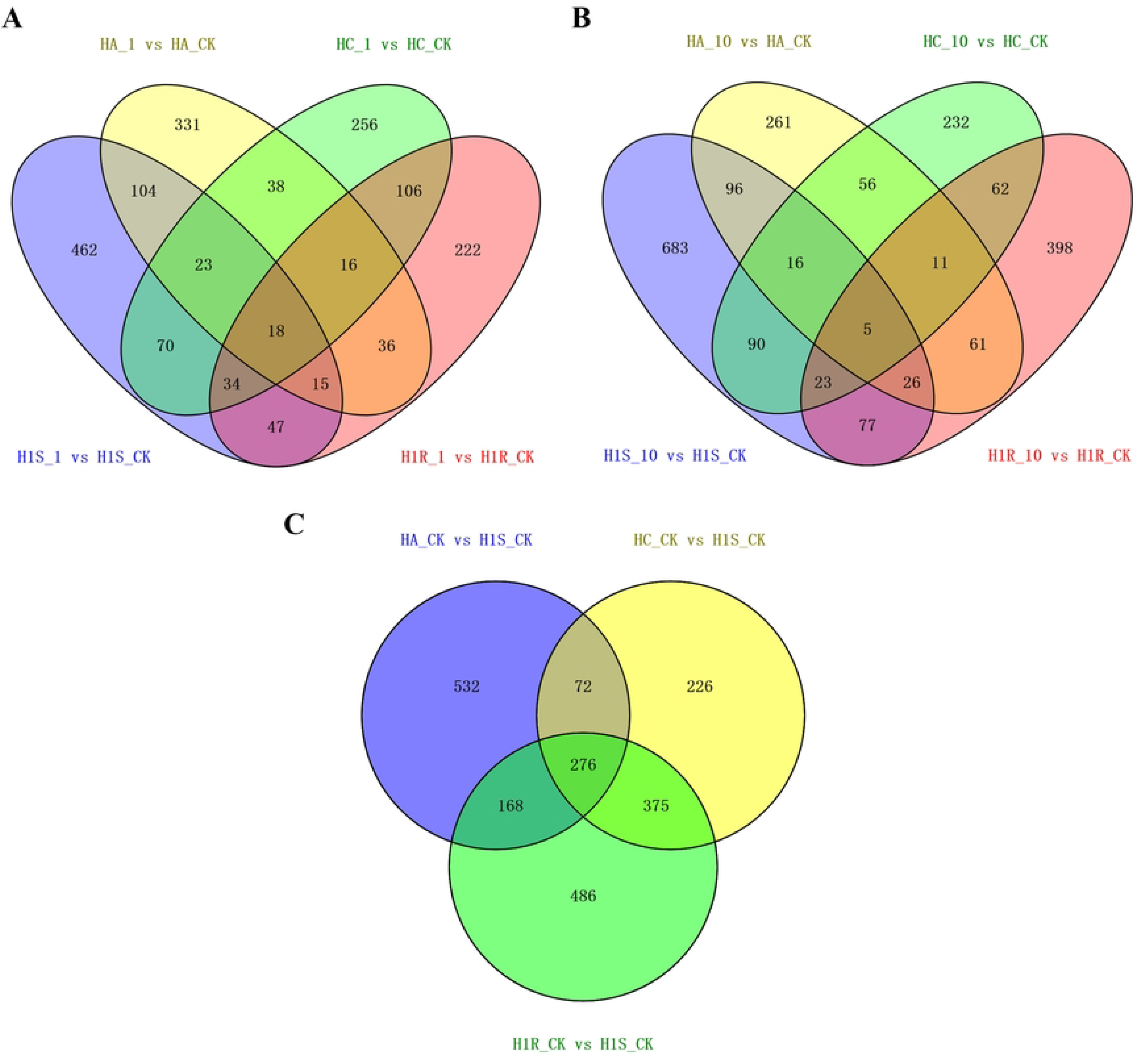
Venn map analysis of DEGs in *F. asiaticum*. (A) Venn plot of DEGs under 1 μg·mL^-1^ phenamacril treatment; (B) Venn plot of DEGs under 10 μg·mL^-1^ phenamacril treatment; (C) Venn map of DEGs between phenamacril-resistant and -sensitive strains.

### GO classification of DEGs after phenamacril treatment

In order to classify the functions of the predicted DEGs in *F. asiaticum*, We performed GO enrichment analysis, which is an internationally standardised gene functional classification system in biological process, cellular component and molecular function. And we chose 30 functional terms based on the significant degree of enrichment analysis from high to low for study. Compared with the phenamacril-sensitive strain H1S, the DEGs of the resistant strains HA, HC, and H1R were significantly co-enriched in small molecule metabolism process, organic acid metabolic process, oxoacid metabolic process, carboxylic acid metabolic process and cellular iron ion homeostasis (FDR<0.05) (Fig 6).

**Fig 6.**
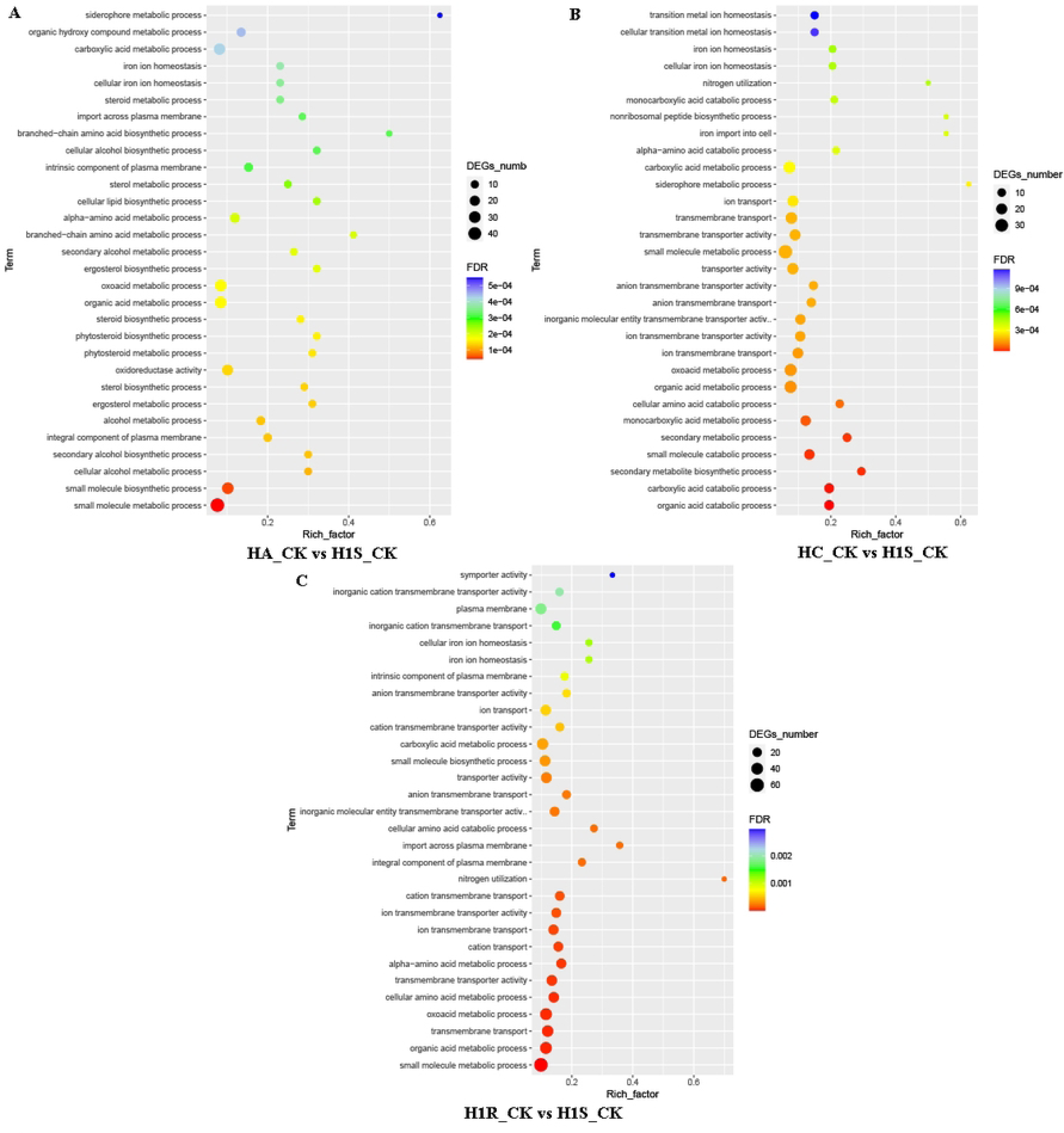
GO enrichment analysis of DEGs between phenamacril-resistant and -sensitive strains comparison groups. The top 30 most enriched GO categories were shown in a scatter diagram in the comparison groups (A) HA_CK vs H1S_CK; (B) HC_CK vs H1S_CK; (C) H1R_CK vs H1S_CK. The abscissa represents the enrichment factor, which is the ratio of the number of DEGs enriched in a certain GO term to the number of background genes obtained through sequencing; The ordinate represents the functions enriched by the GO term. The size of small dots represents the number of genes annotated to GO term, the color from red to purple represents the significance of enrichment, and adjusted FDR < 0.05 is used as the threshold of significance enrichment for GO enrichment analysis.

When treated with 1 μg·mL^-1^ phenamacril, In total 28 and 32 GO terms could be assigned to phenamacril-resistant strains HC and H1R, respectively. In particular, The DEGs of HC_1 vs HC_CK and H1R_1 vs H1R_CK groups (CK represents the control groups) were significantly co-enriched in these GO terms, including cation transmembrane transport, cation transport, ion transmembrane transport, ion transmembrane transporter activity, cation transmembrane transporter activity, organic acid metabolic process, transmembrane transporter activity, oxoacid metabolic process, ion transport, inorganic molecular entity transmembrane transporter activity, transmembrane transport, aromatic amino acid family metabolic process, small molecule metabolic process, inorganic cation transmembrane transporter activity, carboxylic acid metabolic process, secondary active transmembrane transporter activity, cellular amino acid metabolic process, anion transmembrane transporter activity and active ion transmembrane transporter activity (FDR<0.05) (Fig 7B, 7C). There was no enriched GO terms at a threshold of FDR<0.05 but there were 7 enriched GO terms at a threshold of Pvalue <0.05 in the HA_1 vs HA_CK group. These GO terms were involved in small molecule catabolic process, organic substance catabolic process, carbohydrate metabolic process, catabolic process, response to chemical, oxidoreductase activity and catalytic activity (see supporting information Fig S1). However, there were 21 enriched GO terms in the H1S_1 vs H1S_CK group and these GO terms involved in integral component of plasma membrane, intrinsic component of plasma membrane, carboxylic acid catabolic process, import into cell, drug transmembrane transport, xenobiotic transmembrane transporter activity and cellular iron ion homeostasis were only found in the phenamacril-sensitive strains H1S (FDR<0.05) (Fig 7A).

**Fig 7.**
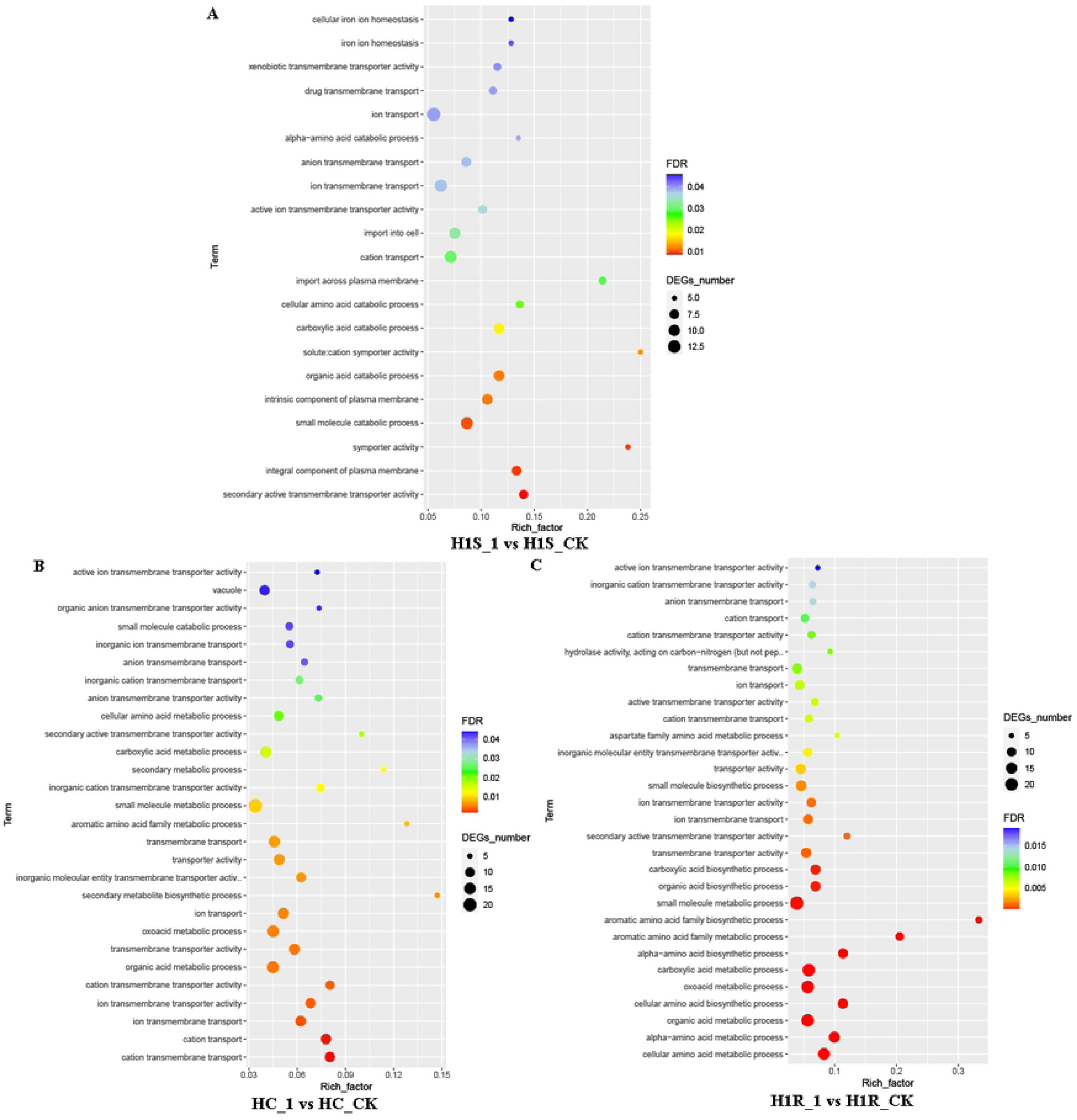
GO enrichment analysis of DEGs in *F. asiaticum* after 1 μg·mL^-1^ phenamacril treatment. The enriched GO categories were shown in a scatter diagram in the comparison groups (A) H1S_1 vs H1S_CK; (B)HC_1 vs HC_CK; (C) H1R_1 vs H1R_CK. The abscissa represents the enrichment factor, which is the ratio of the number of DEGs enriched in a certain GO term to the number of background genes obtained through sequencing; The ordinate represents the functions enriched by the GO term. The size of small dots represents the number of genes annotated to GO term, the color from red to purple represents the significance of enrichment, and adjusted FDR < 0.05 is used as the threshold of significance enrichment for GO enrichment analysis.

When treated with 10 μg·mL^-1^ phenamacril, In total 26 and 21 GO terms could be assigned to phenamacril-resistant strains HC and H1R, respectively. And no DEGs of HC_1 vs HC_CK and H1R_1 vs H1R_CK groups were significantly co-enriched in these GO terms (FDR<0.05) (Fig 8B, 8C). In addition, There was no enriched GO terms at a threshold of FDR<0.05 but there were 19 enriched GO terms at a threshold of Pvalue <0.05 in the HA_1 vs HA_CK group. These GO terms were involved in histone and covalent chromatin modification; DNA, nucleic acid, organic cyclic compound, double-stranded DNA and sequence-specific DNA binding; positive regulation of biosynthetic process, cellular biosynthetic process, macromolecule biosynthetic process, nucleobase-containing compound metabolic process, nitrogen compound metabolic process, gene expression, cellular metabolic process and negative regulation of macromolecule biosynthetic process (see supporting information Fig S2). These results indicated that different types of point mutations lead to different resistance regulation patterns. However, there were 14 enriched GO terms in the H1S_1 vs H1S_CK group and these GO terms involved in organic acid, carboxylic acid, small molecule, antibiotic, monocarboxylic acid, cellular amino acid, organic substance catabolic process and integral component of plasma membrane were only found in the phenamacril-sensitive strains H1S (FDR<0.05) (Fig 8A).

However, KEGG enrichment analysis were not described in this text after phenamacril treatment, and the enriched pathway and data are shown in supporting information Fig S3, Fig S43 and Fig S5.

**Fig 8.**
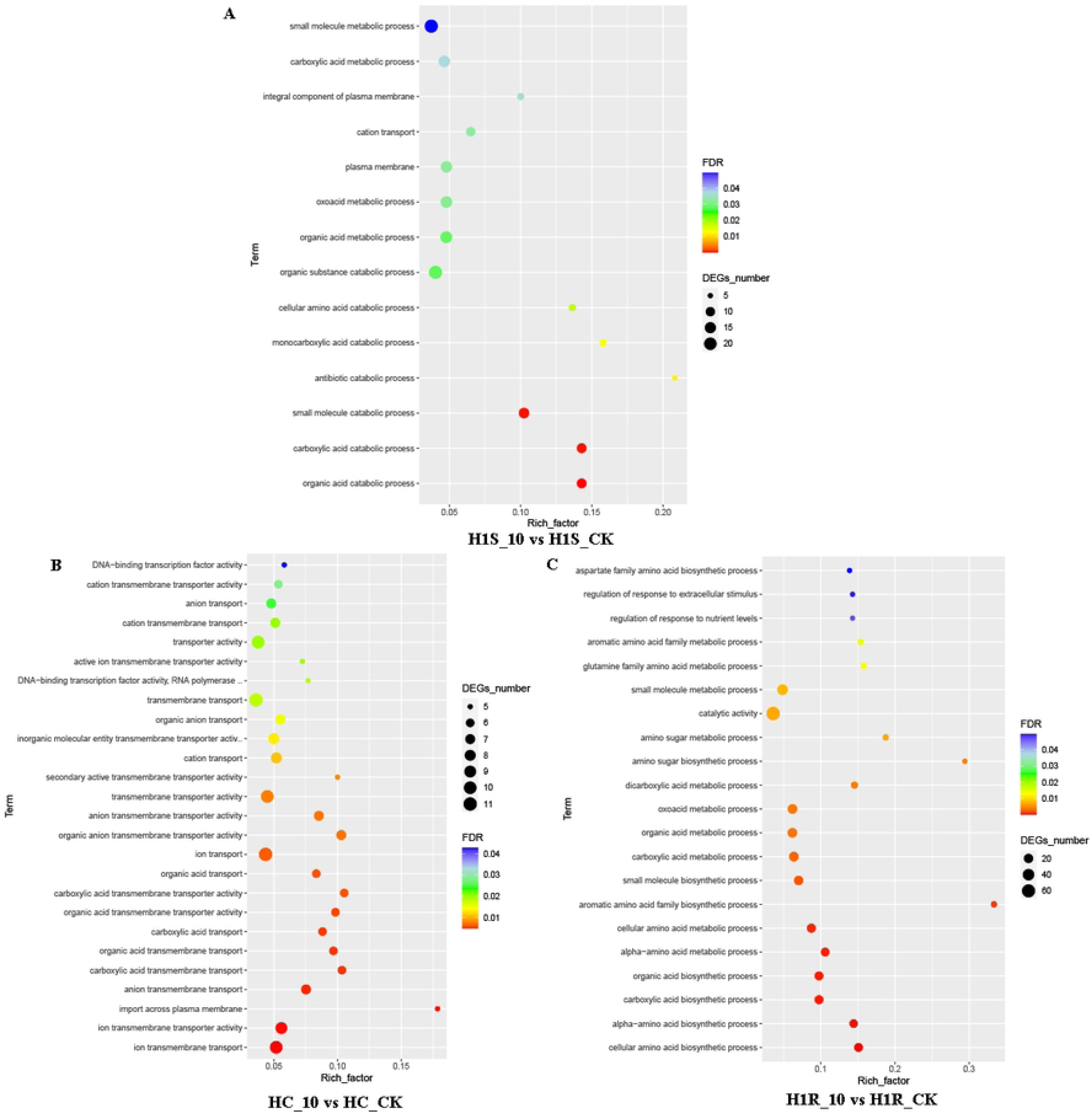
GO enrichment analysis of DEGs in *F. asiaticum* after 10 μg·mL^-1^ phenamacril treatment. The enriched GO categories were shown in a scatter diagram in the comparison groups (A) H1S_10 vs H1S_CK; (B)HC_10 vs HC_CK; (C) H1R_10 vs H1R_CK. The abscissa represents the enrichment factor, which is the ratio of the number of DEGs enriched in a certain GO term to the number of background genes obtained through sequencing; The ordinate represents the functions enriched by the GO term. The size of small dots represents the number of genes annotated to GO term, the color from red to purple represents the significance of enrichment, and adjusted FDR < 0.05 is used as the threshold of significance enrichment for GO enrichment analysis.

### DEGs analysis related to resistance regulation of phenamacril in *F. asiaticum*

In order to screen DEGs related to resistance regulation process, we compared the transcriptomes of phenamacril-resistant strains HA, HC, and H1R with phenamacril-sensitive strain H1S under conditions of 1 μg·mL^-1^, 10 μg·mL^-1^ and no phenamacril treatment. By analyzing the DEGs in the Venn diagram, we listed the selected DEGs after removing false positive expression and extreme repetitive expression. When compared with the phenamacril-sensitive strain H1S, 276 DEGs were co-expressed in the phenamacril-resistant strains HA, HC, and H1R under the condition of no phenamacril treatment. After excluding hypothetical proteins, 15 annotated DEGs were significantly up-regulated and 4 annotated DEGs were significantly down-regulated. The 15 up-regulated DEGs encoded ammonium transporters MEP1/MEP2, methylmalonic-semialdehyde dehydrogenase, pyrimidine precursor biosynthesis enzyme THI12, nitrate reductase, copper amine oxidase 1, glycine cleavage system H protein, amino-acid permease inda1, diphtheria amide biosynthesis protein 3, beta-glucosidase 1 precursor, succinic-semialdehyde dehydrogenase, endothiapepsin precursor, 2,3-dihydroxybenzoic acid decarboxylase and 4-aminobutyrate aminotransferase. The 4 down-regulated DEGs encoded siderophore iron transporter mirB, phenol-2 monooxygenase, iron transport multicopper oxidase FET3 precursor and cysteine dioxygenase (Table 3).

**Table 3.** Expression patterns of sensitive and point mutantion resistant strains of *F. asiaticum*.

| Gene name | Fold change |  |  | Annotation |
| --- | --- | --- | --- | --- |
|  | HA_CK vs | HC_CK vs | H1R_CK vs |  |

|  | H1S_CK | H1S_CK | H1S_CK |  |
| --- | --- | --- | --- | --- |
| FGSG_00490 | 2.34 | 2.26 | 2.20 | methylmalonate-semialdehyde dehydrogenase |
| FGSG_00529 | 2.08 | 2.04 | 3.15 | ammonium transporter MEP1 |
| FGSG_00620 | 3.49 | 6.84 | 8.21 | ammonium transporter MEP2 |
| FGSG_01672 | 2.95 | 4.05 | 5.39 | pyrimidine precursor biosynthesis enzyme THI12 |
| FGSG_01947 | 3.93 | 3.04 | 4.08 | nitrate reductase |
| FGSG_02279 | 7.69 | 2.27 | 2.29 | copper amine oxidase 1 |
| FGSG_03412 | 6.70 | 10.27 | 19.56 | amino-acid permease inda1 |
| FGSG_05554 | 3.58 | 3.83 | 4.88 | 4-aminobutyrate aminotransferase |
| FGSG_06011 | 2.44 | 2.91 | 2.19 | diphthamide biosynthesis protein 3 |
| FGSG_06605 | 3.18 | 3.12 | 4.92 | beta-glucosidase 1 precursor |
| FGSG_06751 | 4.21 | 7.42 | 8.13 | 4-aminobutyrate aminotransferase |
| FGSG_06752 | 16.72 | 24.15 | 25.28 | succinate-semialdehyde dehydrogenase |
| FGSG_07775 | 4.10 | 13.62 | 21.54 | endothiapepsin precursor |
| FGSG_08351 | 3.65 | 2.34 | 2.62 | glycine cleavage system H protein |
| FGSG_09061 | 33.92 | 57.89 | 34.94 | 2,3-dihydroxybenzoic acid decarboxylase |
| FGSG_00539 | -8.34 | -3.12 | -4.31 | siderophore iron transporter mirB |
| FGSG_03199 | -3.67 | -3.21 | -6.13 | phenol 2-monooxygenase |
| FGSG_05159 | -4.91 | -3.92 | -7.17 | iron transport multicopper oxidase FET3 precursor |
| FGSG_09820 | -2.14 | -6.36 | -7.12 | cysteine dioxygenase |

“-” indicates down-regulated expression, “CK” represents control group, fold change ≥ 2 are considered significant differentially expressed genes.

When treated with 1 μg·mL^-1^ phenamacril, 16 genes were significantly expressed in the resistant strains HA, HC and H1R but not in the sensitive strain H1S; These 16 genes encoded hypothetical proteins in *F. asiaticum* and there were 7 genes with functional annotation in other Fusarium species through NCBI BLAST. We found that the DEGs encoded putative Rossmann fold NAD(P)^+^ binding protein (FGSG_10444, FGSG_11100), Medium chain dehydrogenases/zinc alcohol dehydrogenase (FGSG_03913), FAD-binding protein (FGSG_05724), Ig group 2 domain-containing protein (FGSG_11206) and three hypothetical proteins (FGSG_13754, FGSG_08317, FGSG_04021) were significantly down-regulated expression in the phenamacril-resistant strains. On the other hand, there were 462 genes were significantly expressed in the sensitive strain H1S but not in the resistant strains HA, HC and H1R; Among these 462 genes, we counted 21 DEGs with functional annotation in the sensitive strain H1S, of which 13 DEGs were down regulated and 8 DEGs were up regulated. The 8 up-regulated DEGs encoded general stress protein 39, phosphatase 1 regulatory subunit SDS22, copper diamine oxidase, nitrite reductase, glutamine amidotransferase subunit pdxT, drug resistance protein, cytochrome P450 51 protein and NADH dehydrogenase iron-sulfur protein 7; However, the 13 down-regulated DEGs encoded UDP-N-acetylglucosamine pyrophosphorylase, lysophospholipase 2, Ras-2 protein, chitin synthase 1, multiprotein-bridging factor 1, peptidyl-prolyl cis-trans isomerase D, glutathione-independent formaldehyde dehydrogenase, NAD-specific glutamate dehydrogenase, amino-acid permease inda1, betaine aldehyde dehydrogenase, formate dehydrogenase, ATP-dependent RNA helicase DED1 and 4-aminobutyrate aminotransferase (Table 4). When treated with 10 μg·mL^-1^ phenamacril, 11 genes were significantly expressed in the resistant strains HA, HC and H1R but not in the sensitive strain H1S; These 11 genes encoded hypothetical proteins in *F. asiaticum* and there were 4 genes with functional annotation in other Fusarium species through NCBI BLAST.

**Table 4.**
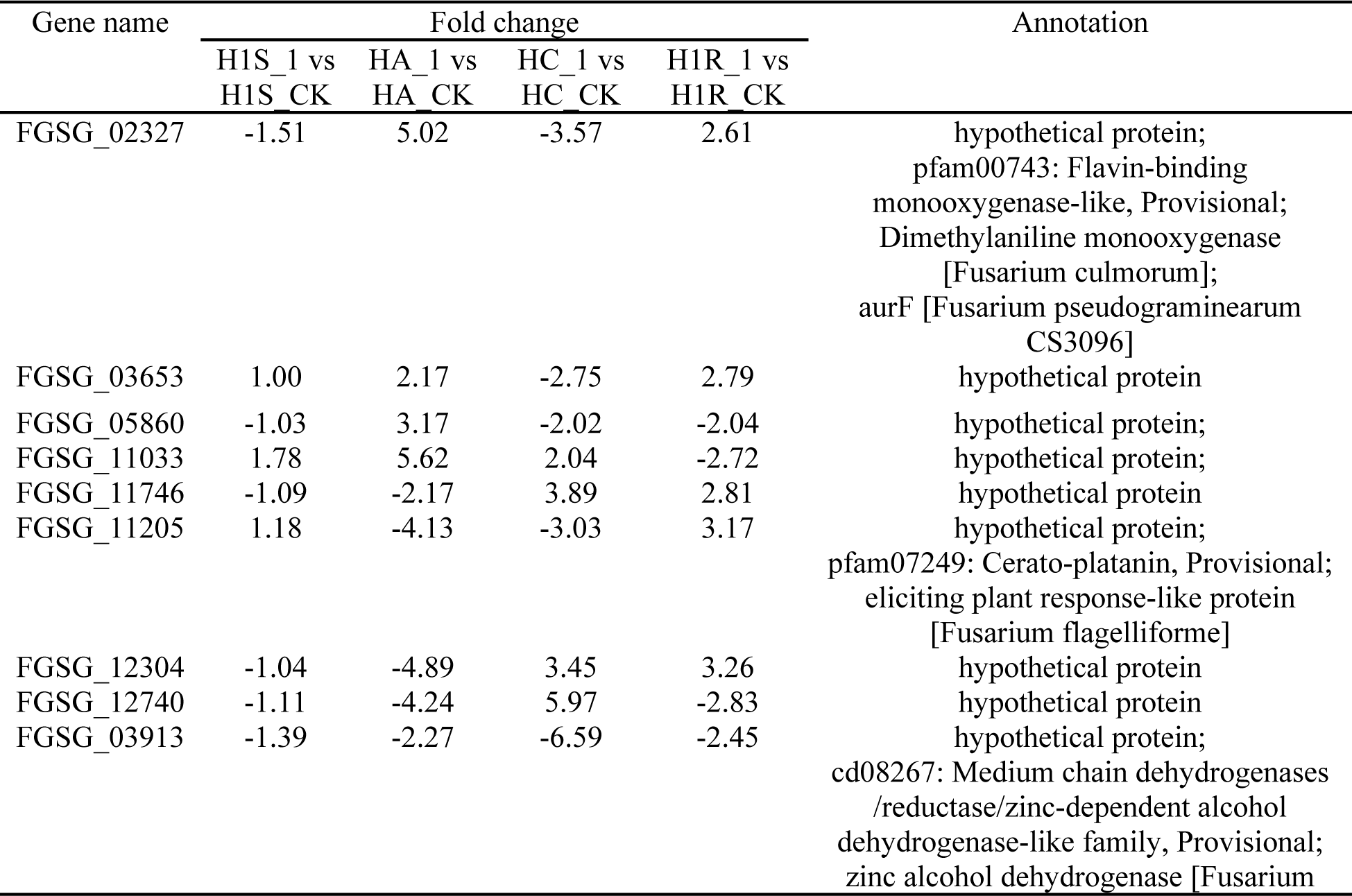
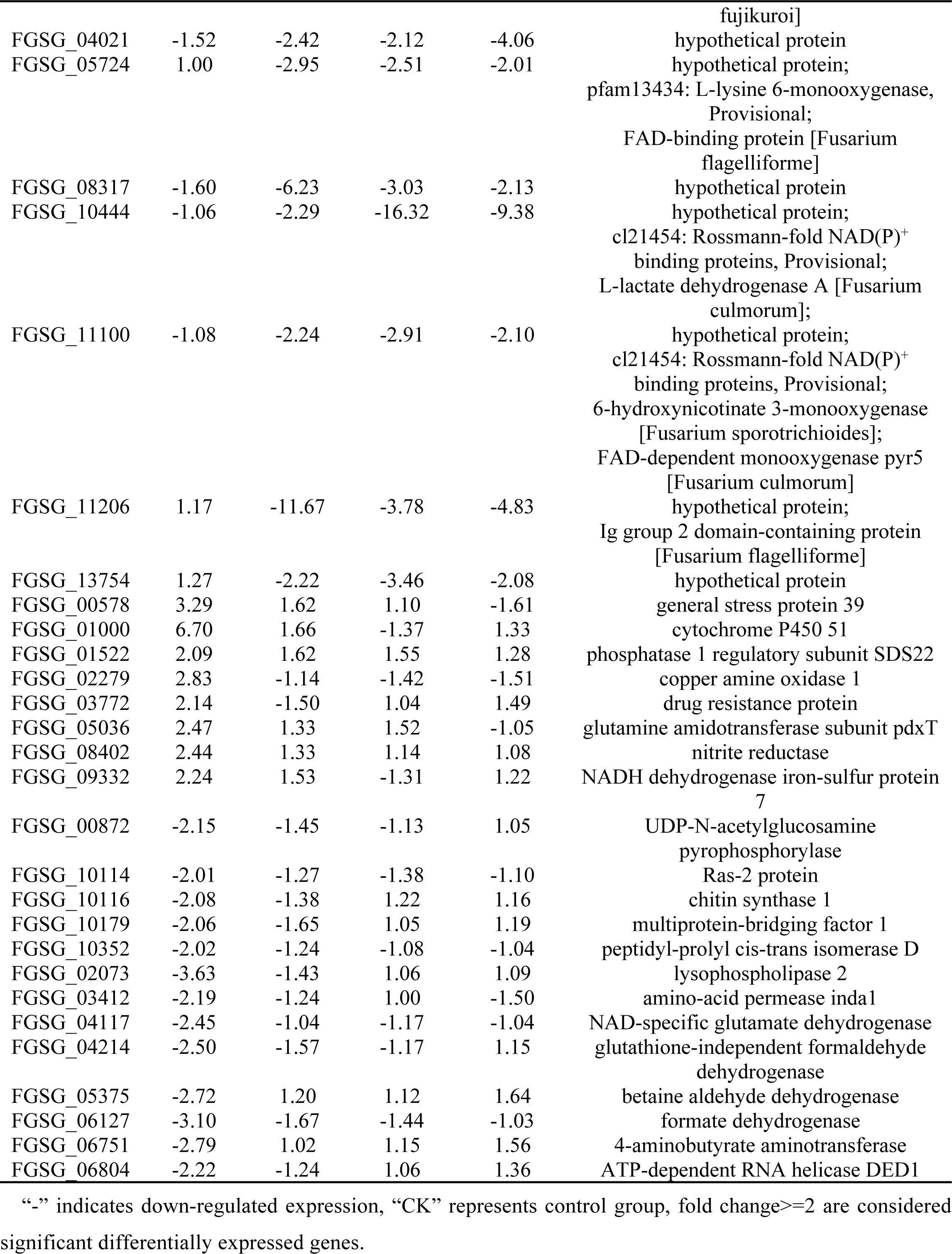
Expression patterns of sensitive and point mutantion resistant strains of *F. asiaticum* under the treatment of 1 μg·mL^-1^ phenamacril.

We found that the DEGs encoded six hypothetical proteins (FGSG_01546, FGSG_11933, FGSG_12185, FGSG_03104, FGSG_13020, FGSG_13022) were significantly down-regulated expression in the resistant strains HA, HC and H1R. On the other hand, there were 683 genes were significantly expressed in the sensitive strain H1S but not in the resistant strains HA, HC and H1R; Among these 683 genes, we counted 27 DEGs with functional annotation in the sensitive strain H1S, of which 14 DEGs were down regulated and 13 DEGs were up regulated. The 13 up-regulated DEGs encoded cytochrome P450 51, P450 61, phosphate carrier protein 2, alcohol dehydrogenase 1, phosphoadenosyl sulfate reductase, glycine dehydrogenase, glycine cutting system H protein, peroxidase/catalase 2, vesicular ATP synthase subunit c, glucan endo-1,3-beta-glucosidase, 5-methyltetrahydropteroyltriglutamate-homocysteine methyltransferase, peroxiredoxin PRX1 and N-amino acid transport system protein. In addition, the 14 down-regulated DEGs encoded acetyl-coenzyme A synthetase, sarcoplasmic/endoplasmic reticulum calcium ATPase 2, ankyrin repeat protein nuc-2, alpha-1,3-mannosyltransferase alg-2, isocitrate lyase, ubiquitin, aldehyde dehydrogenase, import inner membrane translocase subunit TIM22, formate dehydrogenase, glutathione-dependent formaldehyde-activating enzyme, Delta (14)-sterol reductase, S-formylglutathione hydrolase, transcriptional regulatory protein pro-1 and 4-aminobutyrate aminotransferase (Table 5).These genes with different expression patterns in susceptible and resistant strains could explain the mechanism of resistance regulation.

**Table 5.** Expression patterns of sensitive and point mutantion resistant strains of *F. asiaticum* under the treatment of 10 μg·mL^-1^ phenamacril.

| Gene name | Fold change |  |  |  | Annotation |
| --- | --- | --- | --- | --- | --- |
|  | H1S_10 vs<br>H1S_CK | HA_10 vs<br>HA_CK | HC_10 vs<br>HC_CK | H1R_10 vs<br>H1R_CK |  |
| FGSG_00048 | NA | 7.07 | -4.95 | -6.79 | hypothetical protein;<br>COG3491: Isopenicillin N<br>synthase and related<br>dioxygenases, Provisional;<br>putative 2-oxoglutarate-<br>dependent dioxygenase<br>[Fusarium culmorum] |
| FGSG_00049 | NA | 5.09 | -4.24 | -5.38 | hypothetical protein;<br>PRK13357: branched-chain<br>amino acid aminotransferase,<br>Provisional;<br>Aminotransferase apf4 [Fusarium<br>culmorum] |
| FGSG_11500 | 1.81 | 2.57 | -3.37 | -7.02 | hypothetical protein;<br>cl29141 : pyridoxine<br>(pyridoxamine) 5'-phosphate<br>oxidase, Provisional;<br>Pyridoxine/pyridoxamine 5'-<br>phosphate oxidase [Fusarium<br>culmorum] |
| FGSG_12719 | -1.78 | -8.92 | 3.16 | 6.16 | hypothetical protein |
| FGSG_03460 | -1.77 | 2.17 | -2.68 | -5.41 | hypothetical protein;<br>pfam03169 : OPT oligopeptide<br>transporter protein, Provisional;<br>Sexual differentiation process<br>protein isp4 [Fusarium<br>culmorum] |
| FGSG_01546 | -1.31 | -2.54 | -2.44 | -2.08 | hypothetical protein; |
| FGSG_03104 | -1.41 | -2.40 | -5.00 | -2.33 | hypothetical protein |
| FGSG_11933 | 1.10 | -3.30 | -11.75 | -9.35 | hypothetical protein |
| FGSG_12185 | 1.91 | -3.39 | -3.46 | -7.21 | hypothetical protein |
| FGSG_13020 | -1.52 | -3.68 | -3.99 | -2.48 | hypothetical protein |
| FGSG_13022 | 1.14 | -5.39 | -3.36 | -6.21 | hypothetical protein |
| FGSG_01000 | 2.01 | 1.13 | -1.29 | 1.21 | cytochrome P450 51 |
| FGSG_01230 | 2.54 | 1.09 | 1.14 | 1.11 | phosphate carrier protein 2 |
| FGSG_02034 | 5.13 | 1.47 | -1.27 | -1.59 | alcohol dehydrogenase 1 |
| FGSG_03686 | 2.70 | -1.60 | -1.43 | -1.27 | cytochrome P450 61 |
| FGSG_05470 | 4.53 | 1.05 | 1.91 | 1.14 | vacuolar ATP synthase subunit c |
| FGSG_07536 | 2.07 | -1.13 | 1.10 | 1.27 | peroxiredoxin PRX1 |
| FGSG_08351 | 3.57 | -1.15 | -1.24 | -1.19 | glycine cleavage system H<br>protein |
| FGSG_08352 | 3.34 | -1.14 | 1.00 | -1.13 | glycine dehydrogenase |
| FGSG_08528 | 2.04 | -1.29 | 1.63 | 1.67 | phosphoadenosine<br>phosphosulfate reductase |
| FGSG_09354 | 2.16 | -1.20 | 1.63 | 1.40 | N amino acid transport system<br>protein |
| FGSG_10825 | 2.05 | 1.08 | -1.18 | -1.14 | 5-methyltetrahydropteroyltriglutam<br>ate-homocysteine<br>methyltransferase |
| FGSG_11006 | 2.18 | 1.41 | 1.11 | 1.08 | glucan endo-1,3-beta-glucosidase |
| FGSG_12369 | 3.26 | -1.20 | 1.67 | -1.61 | peroxidase/catalase 2 |
| FGSG_00330 | -2.39 | 1.22 | -1.51 | -1.12 | acetyl-coenzyme A synthetase |
| FGSG_01265 | -2.02 | 1.01 | 1.04 | -1.30 | sarcoplasmic/endoplasmic<br>reticulum calcium ATPase 2 |
| FGSG_01544 | -2.24 | -1.08 | -1.18 | 1.27 | ankyrin repeat protein nuc-2 |
| FGSG_01640 | -2.04 | -1.25 | 1.56 | 1.18 | alpha-1,3-mannosyltransferase<br>alg-2 |
| FGSG_05831 | -2.46 | 1.14 | 1.11 | 1.06 | aldehyde dehydrogenase |
| FGSG_06120 | -2.15 | -1.22 | 1.00 | -1.28 | import inner membrane<br>translocase subunit TIM22 |
| FGSG_06127 | -6.28 | 1.01 | -1.17 | -1.07 | formate dehydrogenase |
| FGSG_06606 | -2.10 | 1.15 | -1.46 | 1.06 | Delta (14)-sterol reductase |
| FGSG_06681 | -4.29 | -1.15 | -1.29 | -1.10 | S-formylglutathione hydrolase |
| FGSG_06751 | -2.57 | -1.25 | 1.29 | -1.31 | 4-aminobutyrate<br>aminotransferase |
| FGSG_07067 | -2.43 | 1.02 | -1.30 | -1.29 | transcriptional regulatory protein<br>pro-1 |
| FGSG_08768 | -2.06 | 1.10 | 1.20 | 1.01 | ubiquitin |
| FGSG_09896 | -3.70 | -1.79 | -1.11 | 1.24 | isocitrate lyase |
| FGSG_10776 | -2.00 | -1.07 | 1.17 | 1.17 | glutathione-dependent<br>formaldehyde-activating enzyme |

“-” indicates down-regulated expression, “CK” represents control group, fold change $\geq 2$ are considered significant differentially expressed genes, and NA indicates no detected expression level.

### DEGs analysis related to inhibitory effect of phenamacril against *F. asiaticum*

In order to investigate the inhibitory effect of phenamacril against *F. asiaticum*, We analyzed co-expressed DEGs of phenamacril-resistant strains HA, HC, and H1R and phenamacril-sensitive strain H1S under conditions of 1 μg·mL^-1^ or 10 μg·mL^-1^. The Venn diagram showed that 18 DEGs were co-expressed in both phenamacril-resistant and -sensitive strains when treated with 1 μg·mL^-1^ phenamacril, of which 15 DEGs encoded hypothetical proteins and 3 encoded known proteins. We found that there were 9 genes with functional annotation in other Fusarium species through NCBI BLAST. Interestingly, 14 DEGs were significantly down-regulated in all phenamacril-resistant and -sensitive strains. And these DEGs were involved in putative multidrug resistance protein (FGSG_00541), NADPH-dependent FMN reductase (FGSG_01403), plasma membrane protein yro2 (FGSG_01440), eugenol synthase 1 (FGSG_10433), mating-type protein MAT-1 (FGSG_08892), cyanide hydratase (FGSG_05805), 5-methylthioadenosine s-adenosylhomocysteine deaminase (FGSG_11158), urea amidolyase (FGSG_10913), OPT oligopeptide transporter protein (FGSG_06562), beta-glucosidase 1 precursor (FGSG_06605), purine nucleoside permease (FGSG_07519) and the hypothetical proteins (FGSG_10441, FGSG_10443, FGSG_03970) (Table 6).

**Table 6.**
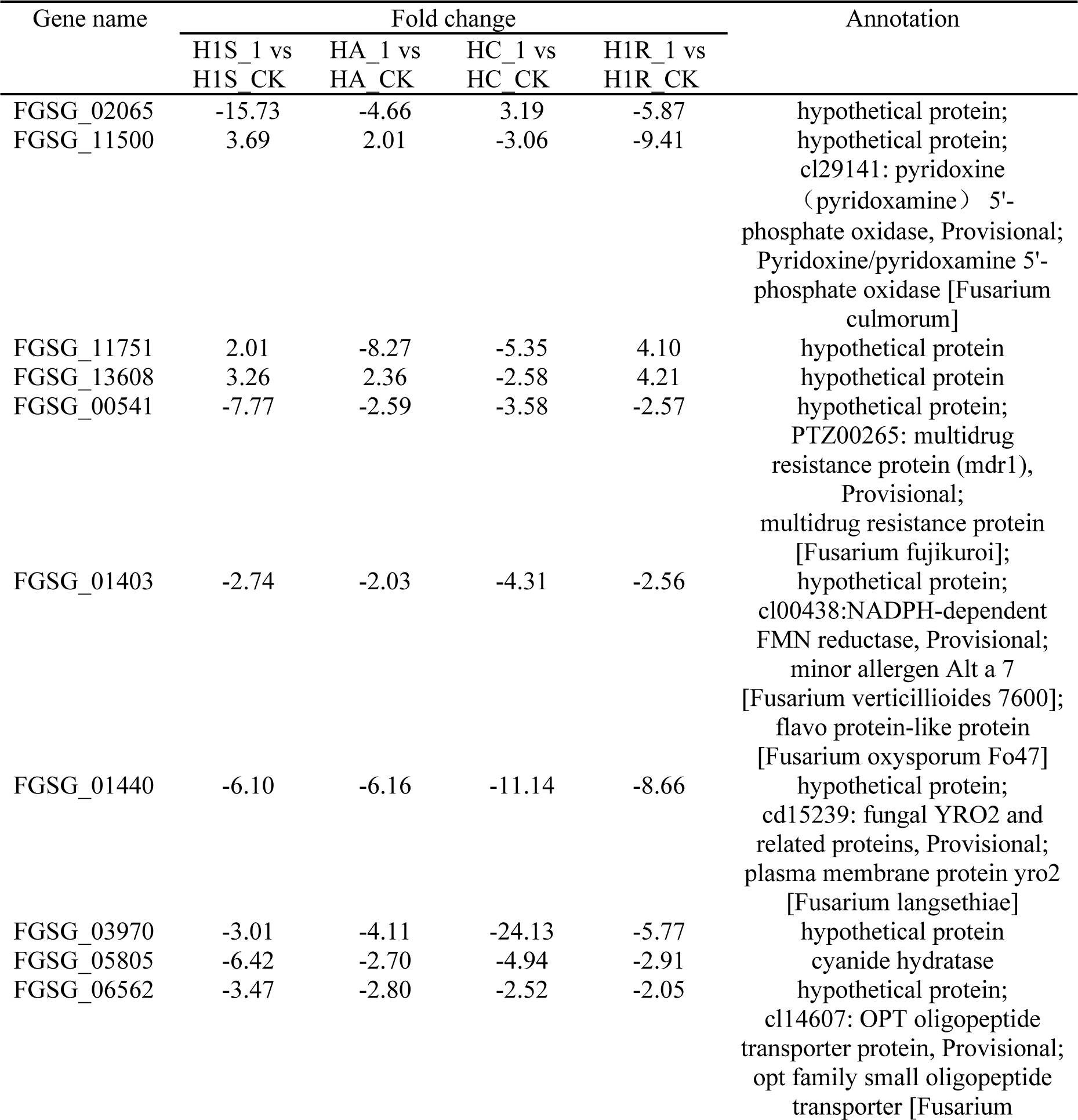
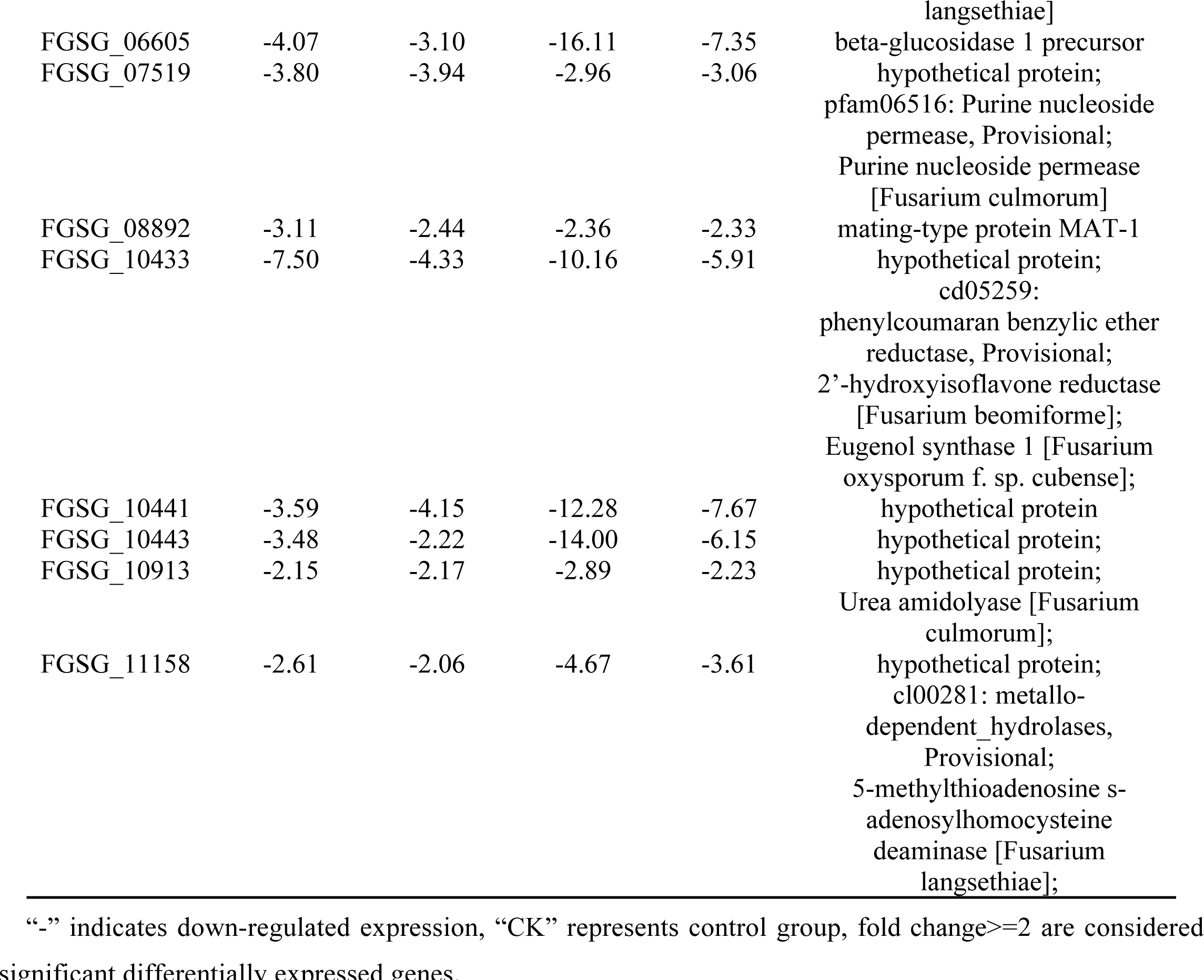
Co-expression patterns of sensitive and point mutantion resistant strainss of *F. asiaticum* under the treatment of 1 μg·mL^-1^ phenamacril.

When treated with 10 μg·mL^-1^ phenamacril, only five DEGs were co-expressed in both phenamacril-resistant and -sensitive strains and all of them encoded hypothetical proteins. We found that two DEGs were significantly down-regulated in all phenamacril-resistant and -sensitive strains. One DEG encodes hypothetical protein (FGSG_06144), another encodes a plasma membrane protein yro2 (FGSG_01440), which was the same as the selected gene treated with 1 μg·mL^-1^ phenamacril (Table 9). In addition, The FGSG_01440 was less down-regulated expression in all the phenamacril-resistant strains when treated with 10 μg·mL^-1^ phenamacril than 1 μg·mL^-1^ phenamacril (Table 7). These results show that phenamacril inhibits these genes in both susceptible and resistant strains.

**Table 7.** Co-expression patterns of sensitive and point mutantion resistant strains of *F. asiaticum* under the treatment of 10 μg·mL^-1^ phenamacril.

| Gene name | Fold change |  |  |  | Annotation |
| --- | --- | --- | --- | --- | --- |
|  | H1S_10 vs<br>H1S CK | HA_10 vs<br>HA CK | HC_10 vs<br>HC CK | H1R_10 vs<br>H1R CK |  |
| FGSG_04142 | -4.31 | -3.74 | 2.02 | 2.05 | hypothetical protein |
| FGSG_08396 | -3.54 | -2.21 | 2.56 | -2.27 | hypothetical protein; |
|  |  |  |  |  | cd00854: N- |
|  |  |  |  |  | acetylglucosamine-6- |
|  |  |  |  |  | phosphate deacetylase, |
|  |  |  |  |  | Provisional; |
|  |  |  |  |  | N-acetylglucosamine-6- |
|  |  |  |  |  | phosphate deacetylase |
|  |  |  |  |  | [Fusarium culmorum] |
| FGSG_13751 | 6.73 | -8.49 | -3.35 | -7.14 | hypothetical protein |
| FGSG_01440 | -7.68 | -2.21 | -5.86 | -3.90 | hypothetical protein; |
|  |  |  |  |  | cd15239: fungal YRO2 and |
|  |  |  |  |  | related proteins, |
|  |  |  |  |  | Provisional; |
|  |  |  |  |  | plasma membrane protein |
|  |  |  |  |  | yro2 [Fusarium langsethiae] |
| FGSG_06144 | -2.11 | -4.28 | -2.04 | -3.41 | hypothetical protein |

“-” indicates down-regulated expression, “CK” represents control group, fold change ≥ 2 are considered

## Disscusion

Phenamacril is a Fusarium-specific cyanoacrylate fungicide used for control of FHB and Rice bakanae disease. In recent years, the inhibitory activities and resistance mechanism of phenamacril to Fusarium spp. have been extensively reported. Consequently, these studies based on genome sequencing, bioinformatics, and gene manipulation have focused on the modes of action of phenamacril and its corresponding resistance mechanisms. Field resistances have been reported in *F. fujikuroi* and are associated with the mutation types of K218T, S219P/L at Myosin-5 in Zhejiang Province since 2016 [17]. But in *F. graminearum* and *F. asiaticum*, we have not found phenamacril-resistant isolates in the field except for those induced by ultraviolet irradiation and phenamacril in the laboratory. The common mutations identified in highly resistant *F. asiaticum* are K216R/E/N, S217P/L and E420K/G/D in FaMyo5 [16]. When the fungicide phenamacril binds to FaMyo5, the ATPase or motor activity is inhibited by blocking ATP turnover and actin binding, resulting in the inhibition of phenamacril-sensitive strains. Mutations in the *FaMyo5* gene that result in binding site modification consequently have non-attachment between phenamacril and the target protein and thus do not affect ATPase or motor activity for phenamacril-resistant strains. Therefore, the Fungicide Resistance Action Committee (FRAC) have listed fungicide phenamacril as cytoskeleton and motor protein classes B6 (FRAC code list 2023, https://www.frac.info/docs/default-source/publications/frac-code-list/frac-code-list-2023---final.pdf).

In this study, we expand our previous work on target resistance of phenamacril and used comparative transcriptome analysis to increase our understanding of resistance regulation mechanism in phenamacril-sensitive strain H1S and point mutation resistant strains HA, HC and H1R (The types of point mutations are S217P, E420G and K216N, respectively). Compared to the phenamacril-sensitive strain H1S, 276 DEGs were co-expressed in the phenamacril-resistant strains HA, HC, and H1R. We found 15 annotated DEGs were significantly up-regulated and 4 annotated DEGs were significantly down-regulated. The up-regulated DEGs are involved in ammonium transporters MEP1/MEP2, amino acid permease inda1, copper amine oxidase 1, nitrate reductase and other metabolic enzymes. Recent studies shows that nitrate is a major source of nitrogen for most bacteria, fungi, higher plants, and this nutrient limits their growth. The first step in the assimilation of nitrate is the influx of nitrate into cells, and it can occur against an electrochemical potential gradient followed by the catalytic activities of nitrate reductase. In addition, loss-of-function nitrate reductase mutants isolated from several nitrate-assimilating fungi appeared to be unable to accumulate nitrate [31].The high affinity ammonium transporters MEP2 is required for pseudohyphal differentiation under standard conditions, whereas the homologous transporters MEP1 are not. Strains lacking MEP2 have no apparent defects in ammonium uptake, metabolism, or growth, even under the low ammonium concentrations that promote filamentation in *Saccharomyces cerevisiae* [32]. Copper amine oxidase (CAO) can catalyse the oxidation of various amine substrates to their corresponding aldehydes, In prokaryotes and fungi, CAOs allow the organisms to utilize amine substrates as sources of carbon and nitrogen [33]. Fungi can also use amino acids as nitrogen or carbon source and to build blocks for protein synthesis. The uptake of amino acids is mediated by amino acid transport systems [34]. Up to 24 members of the amino acid permease family have been found in the yeast *S. cerevisiae* [35], most of which have been functionally characterized [36]. These studies indicated that phenamacril-resistant strains enhanced the ability of uptaking nitrate, ammonia, Lys amino acids and utilizing amine as carbon and nitrogen source to increase the nutrient for mycelial growth.

In addition, we found that succinic semialdehyde dehydrogenase, methylmalonate-semialdehyde dehydrogenase and two 4-aminobutyrate aminotransferases were significantly up-regulated in all the phenamacril-resistant strains. Research shows 4-aminobutyrate aminotransferase converts 4-aminobutyric acid to succinic semialdehyde [37], and succinic semialdehyde dehydrogenase catalyzes the conversion of succinic semialdehyde to succinate [38], thus enters the citric acid cycle. Methylmalonate-semialdehyde dehydrogenase belongs to the CoA-dependent aldehyde dehydrogenase subfamily. It catalyzes the NAD-dependent oxidation of methylmalonate semialdehyde to propionyl-CoA via the acylation and deacylation steps. The propionyl-CoA first converts to succinyl-CoA, and then enters the citric acid cycle for complete oxidative decomposition to produce ATP, which regulated conidiation, polarized germination and pathogenesis in *Magnaporthe oryzae* [39,40]. Therefore, we believe that phenamacril-resistant strains generate more ATP during the tricarboxylic acid process to maintain mycelial growth compared with sensitive strains. These up-regulated DEGs also include pyrimidine precursor biosynthesis enzyme THI12, glycine cleavage system H protein, 2,3-dihydroxybenzoic acid decarboxylase, beta-glucosidase 1 precursor, diphtheria amide biosynthesis protein 3 and endothiapepsin precursor. Glycine cleavage system H protein is a component of the glycine cleavage system [41], a conserved protein complex that acts to decarboxylate glycine. 2,3-Dihydroxybenzoic acid decarboxylase, which is the last enzyme in the fungal metabolism of indole to catechol, catalyzes the non-oxidative decarboxylation of 2,3-dihydroxybenzoic acid to catechol [42]. β-Glucosidase is an important component of the cellulase enzyme system. It does not only participate in cellulose degradation, but also plays an important role in hydrolyzing cellulose to fermentable glucose by relieving the inhibition of exoglucanase and endoglucanase from cellobiose [43]. Wang et al., (2016) reported that pyrimidine precursor synthase gene *MrThi12* deletion mutant could not produce aerial hyphae and conidia on the vitamin B1-lacking media in *Metarhizium robertsii* [44]. These studies indicated that phenamacril-resistant strains also had stronger abilities of glycine or indole metabolism, cellulose degradation and uptaking vitamin B1, which ensure that resistant strains can utilize more carbon or nitrogen source, and vitamins to survive better.

The 4 down-regulated DEGs encoded siderophore iron transporter mirB, iron transport multicopper oxidase FET3 precursor, phenol-2 monooxygenase and cysteine dioxygenase. Siderophores have been identified as virulence factors in fungus *Aspergillus fumigatus*. The 14-pass transmembrane protein MirB is postulated to function as a siderophore transporter, responsible for uptake of the hydroxamate siderophore N,N′,N″-triacetylfusarinine C [45]. The multicopper oxidase FET3 of yeast can oxidize extracellular ferrous iron Fe ^2+^ and translocate Fe^3+^ to the permease Ftr1 through a pathway under electrostatic guidance, which is then transported into the cell through the permease Ftr1 [46]. The monooxygenase phenol hydroxylase (PheA) can catalyze the first step in the degradation of phenol in the thermophilic bacterium *B. thermoglucosidasius* A7 when the strain is grown on phenol as carbon source and energy. The phenol-2 monooxygenase (PheA2) is an NADH-dependent FAD-containing reductase that supplies PheA1 with reduced FAD to catalyze the *ortho*-hydroxylation of simple phenols to the corresponding catechols and plays a direct role for the protein-bound FAD in the mediation of electron transfer from NAD(P)H to exogenous free flavin [47]. L-Cysteine is an important precursor for ribosomal and nonribosomal peptide synthesis, a sulfur donor for the construction of iron sulfur cluster in proteins, for the biosynthesis of vitamins and secondary metabolites, the thiolation of nucleic acids, or production of the signaling molecules hydrogen sulfide and polysulfides. For cysteine degradation, one important pathway is catalyzed by cysteine dioxygenases. This enzyme catalyzes the four-electron oxidation of cysteine to cysteine sulfinic acid in a oxygen and iron-dependent reaction [48]. These studies indicated that phenamacril-resistant strains not only reduced iron transport and uptake, but also reduced phenol and cysteine degradation, which save more carbon, nitrogen source and an important precursor L-cysteine to maintain various cellular life activities.

In order to investigate the response of sensitive and resistant strains to phenamacril. We compared the differentially expressed genes between the two types of strains under 1 μg·mL^-1^ and 10 μg·mL^-1^ phenamacril treatment. When treated with 1 μg·mL^-1^ phenamacril, the mycelial growth of sensitive strains was severely inhibited, while the resistant strains were not affected on PDA plates. DEGs analysis showed that putative Rossmann fold NAD(P)^+^ binding protein, Medium chain dehydrogenases/zinc alcohol dehydrogenase, FAD-binding protein, Ig group 2 domain-containing protein and three hypothetical proteins (FGSG_13754, FGSG_08317, FGSG_04021) were significantly down-regulated expression in the phenamacril-resistant strains. The Rossmann fold was first identified in the dinucleotide-binding proteins and the fold that serves as flavin adenine dinucleotide (FAD) and pyridine nucleotide (NAD/NADP) [49]. FAD-binding proteins play a vital role in energy transfer and utilization during fungal growth and mycelia aggregation [50]. RNA binding occurs in the Rossmann fold or the NAD^+^-binding region, which permits a rationale for the RNA binding activity attributed to several dehydrogenases and other enzymes. The enzymes containing this domain may exhibit specific RNA binding activity and play additional roles in nucleic acid metabolism [51]. Alcohol dehydrogenases (ADHs) catalyse the reversible oxidation of alcohols to aldehydes or ketones with the corresponding reduction of NAD^+^ or NADP^+^ and the medium-chain dehydrogenases (MDRs) are the member of these enzymes. MDRs that use zinc in their catalytic reaction possess a sequence motif known as the zinc-containing ADH signature [52]. From these studies we speculated that resistant strain might avoide damage from phenamacril by reducing aldehyde production, decreasing energy transfer and utilization during fungal growth, and interfering nucleic acid metabolism through inhibiting RNA binding activity.

On the other hand, there were up to 462 DEGs expressed in the sensitive strain H1S but not in three resistant strains, of which 8 annotated DEGs were up-regulated and 13 annotated DEGs were down-regulated. We found that these up-regulated DEGs were involved in general stress protein 39, phosphatase 1 regulatory subunit SDS22, copper diamine oxidase, nitrite reductase, glutamine amidotransferase subunit pdxT, drug resistance protein, cytochrome P450 51 protein and NADH dehydrogenase iron-sulfur protein 7. Research shows *Bacillus subtilis* can induce a lot of general stress proteins in response to a salt or heat stress which enabled them to survive [53]. Protein phosphatase type 1 plays a major role in the regulation of glycogen biosynthesis which results in glycogen accumulation [54]. The nitrite reductase can degrade nitrite to NO or NH_3_, thereby reducing the toxic effects of nitrite accumulation on organisms. Glutamine amidotransferases can generate ammonia from the glutamine amide nitrogen and transfer it to seven different chemical classes of acceptors which involved in the biosynthesis of nucleotides, amino acids, aminated sugars, coenzymes and antibiotics [55]. The Cytochrome P450 51 (Cyp51) is required for sterol biosynthesis in different phyla and the overexpression of the *cyp51* gene can increase fungicide resistance [56]. NADH dehydrogenase iron-sulfur protein plays a vital role in cellular ATP production, which is the primary source of energy for many crucial processes in living cells [57]. From these studies we proposed that phenamacril-sensitive strain can maintain basic life activities by inducing stress proteins or drug resistance proteins, reducing the toxic effects, enhancing fungicide metabolism, accumulating glycogen or ammonia and generating more energy in response to 1 μg·mL^-1^ phenamacril stress.

In addition, the 13 down-regulated DEGs were involved in UDP-N-acetylglucosamine pyrophosphorylase, chitin synthase 1, glutathione-independent formaldehyde dehydrogenase, formate dehydrogenase, NAD-specific glutamate dehydrogenase, amino-acid permease inda1, betaine aldehyde dehydrogenase, ATP-dependent RNA helicase DED1, 4-aminobutyrate aminotransferase, Ras-2 protein, multiprotein-bridging factor 1, lysophospholipase 2, and peptidyl-prolyl cis-trans isomerase D. Recent studies indicates that UDP-N-acetylglucosamine pyrophosphorylase (UAP1) catalyses the last step in eukaryotic biosynthesis of uridine diphosphate-N-acetylglucosamine (UDP-GlcNAc), converting UTP and GlcNAc-1P to the sugar nucleotide[58, 59]. Chitin is widely found in the cell walls of filamentous fungi and Chitin synthesis is catalyzed by chitin synthase, an enzyme that transfers GlcNAc from UDP-GlcNAc to the nonreducing end of growing chitin chains [60] (Sanchez-Leon). Formaldehyde dehydrogenase (FDH) is a member of the zinc-containing medium-chain alcohol dehydrogenase family which oxidizes toxic formaldehyde to formate using NAD+ as an electron carrier and most FDHs are dependent on glutathione for catalysis [61]. And the formate dehydrogenases catalyse the oxidation of formate to carbon dioxide (CO_2_), which completely oxidizes and decomposes toxic formaldehyde [62]. Glutamate dehydrogenase (GDH) catalyzes the reversible amination of 2-oxoglutarate with ammonium to form glutamate and plays a role in nitrogen assimilation in *A. nidulans* [63] (Tomohito). The NAD-specific glutamate dehydrogenase of *Neurospora crassa* is regulated by carbon catabolite repression being derepressed during growth on glutamate or simply by carbon starvation [64]. Betaine aldehyde dehydrogenase (BADH) catalyzes the NAD(P)^+^-dependent irreversible oxidation of betaine aldehyde to glycine betaine in varieties of organisms, ranging from bacteria to mammals. Glycine betaine is a compatible osmolyte or osmoprotectant synthesized and accumulated at high concentrations in cells under osmotic stress. Therefore, BADH plays an important role in the water stress response of many organisms [65]. The ATP-dependent RNA helicase DED1 from yeast are critical for translation initiation and the repression of DeD1 activity can lead to accumulation of RNA structure in 5′UTRs, translation initiation from near-cognate start codons immediately upstream of these structures and decreased protein synthesis from the corresponding main ORFs[66]. Multiprotein bridging factor 1 is a highly conserved protein in archaea and eukaryotes, which was demonstrated to enhance transcription by forming a bridge between distinct regulatory DNA-binding proteins and the TATA-box-binding protein [67]. Ras-2 protein is localized to the plasma membrane and plays a role in signal transduction [68]. From these studies we concluded that 1 μg·mL^-1^ phenamacril inhibited the mycelial growth of sensitive strain by interfering with cell wall formation, reducing the rate of succinic semialdehyde generation or toxic formaldehyde decomposition, reducing the ability of repressing carbon starvation, transporting amino acid or response to water stress, decreasing transcription or protein synthesis and obstructing signal transduction, while the resistant strains were not affected.

When the treatment concentration of phenamacril increases to 10 μg·mL^-1^, sensitive strains could not grow, while the mycelial growth of resistant strains were slightly inhibited on PDA plates. DEGs analysis indicated that the co-downregulated expression genes in resistant strains encoded six hypothetical proteins (FGSG_01546, FGSG_03104, FGSG_11933, FGSG_12185, FGSG_13020, FGSG_13022). In addition, there were up to 683 DEGs expressed in the sensitive strain H1S but not in three resistant strains, of which 13 annotated DEGs were up-regulated and 14 annotated DEGs were down-regulated. We found that these up-regulated or down-regulated DEGs were involved in similar proteins or enzymes above. And we believe that 10 μg·mL^-1^ phenamacril further inhibit the survival of sensitive strain by increasing aldehyde production (alcohol dehydrogenase 1) and reducing the degradation of toxic aldehyde and formaldehyde (aldehyde dehydrogenase, formate dehydrogenase), increasing the expression of vacuolar ATP synthase subunit c and reducing the expression of sarcoplasmic/endoplasmic reticulum calcium ATPase 2, enhancing the hydrolysis of fungal cell walls (glucan endo-1,3-beta-glucosidase) [69], decreasing ergosterol, acetyl-CoA biosynthesis and conversion of isocitrate (Delta (14)-sterol reductase, acetyl-coenzyme A synthetase, isocitrate lyase), and affecting other metabolic enzymes. In addition, research shows Sarco/endoplasmic reticulum calcium ATPase enzymes play important roles in several signal transduction pathways that control proliferation, differentiation and apoptosis[70] The multi-subunit vacuolar ATPase pump uses ATP hydrolysis to move protons into membrane bound compartments, which is involved in a variety of cellular functions, including regulation of cytosolic pH, vesicular transport, endocytosis, secretion, and apoptosis [71].Taken together, phenamacril have a greater impact on the integrity of the cell wall and cell membrane of sensitive strains, as well as the transport and metabolism of substances and energy, and the synthesis of nucleic acids and proteins compared to resistant strains.

Finally, we also explored the inhibitory effect of phenamacril by investigating the common inhibitory genes on sensitive and resistant strains. From co-expressed DEGs of sensitive and resistant strains under conditions of 1 μg·mL^-1^ or 10 μg·mL^-1^ phenamacril treatment, we have selected mating-type protein MAT-1, cyanide hydratase, beta-glucosidase 1 precursor, putative multidrug resistance protein, NADPH-dependent FMN reductase, plasma membrane protein yro2, eugenol synthase 1, 5-methylthioadenosine s-adenosylhomocysteine deaminase, urea amidolyase, OPT oligopeptide transporter protein, and purine nucleoside permease. Studies indicates that mating type genes have been characterized in numbers of homothallic, heterothallic, and asexual filamentous *Ascomycetes*. In fertile fungi, the mating-type protein MAT-1 functions as a master regulatory protein controlling sexual reproduction [72]. 1 μg·mL^-1^ phenamacril treatment reduced the expression of mating-type protein MAT-1 in both sensitive and resistant *F. asiatiucm* strains, which can explain why we have not detected any phenamacril-resistant isolates in the fields for so many years. Because ascospores from sexual reproduction of Fusarium *spp.* are responsible for the primary infection of wheat spikes during wheat flowering, the phenamacril-resistant strains are unable to overwinter due to their inability to reproduce sexually, thereby reducing the accumulation of resistant populations in the fields. Multidrug resistance (MDR) is an important resistance mechanism. In fungi, the major types of drug efflux proteins are ATP binding cassette (ABC) and major facilitator superfamily (MFS) transporters. Several efflux transporter genes have been shown to be rapidly induced by fungicides or natural toxins[73]. MDR1 is now recognized to be an important protein that regulates the pharmacokinetics of various types of structurally unrelated drugs by transporting them from the inside to the outside of the cell [74]. NADPH-dependent FMN reductase plays a role in many cellular processes by transferring electrons to a protein substrate or another electron acceptor. These enzymes are involved in essential bioenergetic processes, biochemical degradations, and biosynthesis and detoxification reactions [75]. In *S. cerevisiae*, yro2 have seven predicted transmembrane domains and localize to the plasma membrane. The null mutants of yro2 gene exhibited delayed growth and decreased ability to produce ethanol in the presence of acetic acid, indicating that yro2 is involved in tolerance to acetic acid stress [76]. From these studies we proposed that phenamacril performed the inhibitory effect by interfering with sexual reproduction and energy generation, affecting cell wall synthesis and toxic cyanide hydrolysis, reducing multidrug resistance, weakening tolerance to acetic acid stress, repressing oligopeptide and nucleotide bases transport in *F. asiatiucm*.

In summary, fungicide resistance occur in fungi and the resistance are associated not only with target sites but also with non-target site mechanisms [77,78]. Even though we know the point mutations in the gene encoding FaMyo5 confer a high level of resistance, the resistance regulation mechanisms are complicate due to the presence of metabolic resistance. From our research, it can be seen that the response of sensitive and resistant strains to phenamacril is systematic and complex in *F. asiaticum*, which involving in large numbers of proteins and enzymes. In addition to functional annotated proteins, there are many hypothetical proteins worth for further study. Furthermore, the resistance regulation mechanism and inhibitory effect of phenamacril in *F. asiaticum* we have discovered provides a reference basis for the study of drug resistance in other microorganisms. In addition, the relevant proteins or enzymes annotated in our research also have reference value for the discovery of new fungicide targets.

## Materials and methods

### Fungicide and media

Technical-grade phenamacril (95%, CAS: 39491-78-6) was provided by the Jiangsu Pesticide Research Institute Co., Ltd, Nanjing, China and dissolved in methanol to 4, 10 and 40 mg·mL^-1^ for stock solution. The strains used in this study are listed in Table 1 and included the *F. asiaticum* sensitive strains H1, H1S and high resistant strains HA, HC, H1R, Y2021A, Y2021C and HR. All the strains were routinely maintained at 25 °C on Difco™ Potato Dextrose Agar plates (PDA, suspend 46 g of the powder in 1 L of purified water and autoclave at 121 °C for 20 min). For mycelial growth assays, the strains were grown at 25 °C on PDA plates for 3 days. For conidia production, 15 fresh mycelial plugs taken from the periphery of a 3-day-old colony of *F. asiaticum* phenamacril-sensitive strain H1 were added to a 250 mL flask containing 150 mL of Mung bean liquid (MBL, 30 g of mung bean boiled in 1 liter of water for 20 min; the mixture was then filtered through cheesecloth) medium.

### Construction of point mutation vectors for FaMyo5 and transformation

The 2.3-kb *FaMyo5* gene fragments of *F. asiaticum* were amplified from wild-type strain H1 and the resistant strains HR, Y2021A, Y2021C with primers FaMyo5-downF/FaMyo5-downR (Table 2). The HPH resistance gene containing the *Aspergillus nidulans* trpC promoter was amplified from plasmid pKHt with primers hphF/hpH1R. A 1.2-kb upstream flanking fragment of *FaMyo5*, which is located 166 bp upstream from the start codon, was amplified from strain H1 with primers FaMyo5-upF/FaMyo5-upR (Table 2). The amplicons (upstream junction, trpC + HPH, and downstream junction) were amplified with Vazyme’s Phanta Max Super-Fidelity DNA Polymerase and purified with Omega E.Z.N.A. Gel Extraction Kit (D2500-02, Shanghai, China) for the fusion. Then, we used primer pairs GFP-S/GFP-A to amplify vector fragments from plasmid 1300-*HYG*-*sGFP* and used Vazyme’s ClonExpress MultiS One Step Cloning Kit (Nanjing, China) to connect vector fragments and amplicons. Finally, we transformed the connected plasmid into *Escherichia coli* DH5-α competent cells. After selecting the true positive colony and colony PCR validation, we extracted the targeted plasmids and sequenced in Tsingke Biotechnology Co., Ltd to verify the accuracy of point mutation vectors.

For protoplast transformation, 2 μg of fused fragments were amplified from the correct point mutation vectors with primers FaMyo5F/FaMyo5R. The fused products were used to transform protoplasts of the wild-type strain H1. The protoplast preparation and transformation of *F. asiaticum* were performed as previously described [15]. Putative transformants were selected in the presence of hygromycin B (100 μg/mL) and checked by DNA gel blot analysis.

### Fungicide susceptibility testing

Susceptibility to the fungicide phenamacril was assessed for phenamacril-sensitive strains H1, H1S and phenamacril-resistant strains HA, HC and H1R listed in Table 1. For testing inhibition of mycelia growth, mycelial plugs (5 mm in diameter) taken from the margin of a 3-day-old colony were placed on the center of PDA plates amended with phenamacril at: 0, 0.2, 0.4, 0.8, or 1.6 μg·mL^-1^ for sensitive strains; 0, 50, 100, 200, or 400 μg·mL^-1^ for high resistant strains determined by EC_50_ values as the previous classification methods [16]. Three replicates for each concentration were used for each strain. After cultures were kept at 25 °C for 3 days, the colonies were photographed and colony diameters were measured; the diameter (5 mm) of the original mycelial plugs were subtracted from each measurement. The 50% efective concentration (EC_50_) values of strains were calculated by regressing percentage growth inhibition against the log of fungicide concentration with DPS v9.01 sofware (Hangzhou Reifeng Information Technology Ltd., Hangzhou, China) Each experiment yields a set of EC_50_s and the experiment was performed three times.

### Sequence alignment of FaMyo5 motor domains

All the *F. asiaticum* transformants were cultured in PDB at 25 °C for 2 days and the mycelia were collected and finely ground to a powder using a mortar and pestle with liquid nitrogen. The total RNA was extracted using the E.Z.N.A. Fungal RNA Kit (Omega Bio-tek, Inc., Norcross, USA) following the manufacturer’s instructions and used for reverse transcription with the PrimeScript^TM^ RT reagent Kit (TaKaRa).The sequences of *FaMyo5* motor domains were amplified from the cDNAs of all the *F. asiaticum* transformants using the primer pairs myosin5F/myosin5R. The target fragments were purified and cloned into the PMD18-T vector, then sequenced in Tsingke Biotechnology Co., Ltd. and aligned using Bioedit 7.2 software (Isis Pharmaceuticals).

### Sampling for RNA extraction

The conidia harvested from 7-day-old MBL cultures of one phenamacril-sensitive strain H1S and tH1Ree phenamacril-resistance strains HA, HC and H1R were collected and suspended in sterile distilled water at 1×10^6^ conidia/mL. The freshly harvested conidia of each strain were cultured in tH1Ree flasks containing 100ml liquid YEPD medium (w/v, 1% peptone, 0.3% yeast extract, 2% glucose) on a shaking table at a speed of 175 r/min for 18 h at 25 °C. Then we added 25 μL 4 mg·mL^-1^ and 40 mg·mL^-1^ phenamacril into the flasks of treatment groups and made the final concentration of phenamacril 1 μg·mL^-1^ and 10 μg·mL^-1^. We also added 25 μL methanol into the flasks of control groups (CK) to eliminate the influence of solvent methanol on experimental treatment. A total of 4 control strains and 8 treatment strains continued to be cultured for 12 h at 25 °C. After 30 h, the young mycelium were collected and finely ground to a powder using a mortar and pestle with liquid nitrogen and the total RNA was extracted using the above method. All treatments and controls were performed independently tH1Ree times and finally we got 36 RNA samples.

### RNA-seq libraries construction and Illumina sequencing

Total RNA were extracted from the tissue using TRIzol® Reagent according the manufacturer’s instructions (Invitrogen) and genomic DNA was removed using DNase I (TaKara). Then RNA quality was determined using Agilent 2100 BioAnalyzer (Agilent Technologies, Santa Clara, CA, USA) and quantified using the ND-2000 (NanoDrop Technologies). High-quality RNA sample (OD260/280=1.8∼2.2, OD260/230≥2.0, RIN≥7, 28S:18S≥1.8, >10 μg) is used to construct sequencing library. RNA-seq transcriptome libraries were prepared following the NEB Next Ultra RNA Library Prep Kit from Illumina (NEB). NEB Next Poly(A) mRNA Magnetic Isolation Module (NEB) kit was used to enrich the poly(A) tailed mRNA molecules from 1 μg total RNA. The mRNA was fragmented into 200 base pair pieces. The first-strand cDNA was synthesized from the mRNA fragments reverse transcriptase and random hexamer primers, and then the second-strand cDNA was synthesized using DNA polymerase I and RNaseH. The end of the cDNA fragment was subjected to an end repair process that included the addition of a single “A” base, followed by ligation of the adapters. Products were purified and enriched by polymerase chain reaction (PCR) to amplify the library DNA. The final libraries were quantified an Agilent 2100 Bioanalyzer. After quantitative reverse transcription-polymerase chain reaction (RT-qPCR) validation, paired-end libraries were sequenced by Illumina Novaseq 6000 platform (150bp*2, CapitalBio Technology Co., Ltd, Beijing, China).

### RNA-Seq Data Analysis

The raw paired end reads, which removed the reads containing poly-N using in-house perl scripts, were trimmed by Trimmomatic [79] with parameters (SLIDINGWINDOW:4:15 MINLEN:75) (version 0.36). Then clean reads were separately aligned to *Fusarium graminearum* PH-1 reference genome (https://www.ncbi.nlm.nih.gov/datasets/genome/GCF_000240135.3/) with orientation mode using HISAT2 v2.0.5 (https://daehwankimlab.github.io/hisat2/) software with default parameters. The quality assessment of these data were taken by qualimap_v2.2.1 (http://qualimap.bioinfo.cipf.es/). The mapped reads of each sample were assembled by StringTie (v1.3.3b) in a reference-based approach [80]. FeatureCount v1.6.3 (https://subread.sourceforge.net/) was used to count the reads numbers mapped to each gene [81]. To identify DEGs (differential expression genes) between the two different samples, the expression level for each gene was calculated using the fragments per kilobase of exon per million mapped reads (FRKM) method. R statistical package edgeR3.6.3 (Empirical analysis of Digital Gene Expression in R, http://www.bioconductor.org/packages/release/bioc/html/edgeR.html/) was used for differential expression comparison. DESeq2 was used to analyze the DEGs between samples. The resulting p-values were corrected using the Benjamini and Hochberg’s method for controlling the false discovery rate (FDR). Parameters for classifying significantly DEGs are ≥2-fold differences (|log_2_FC|≥1, FC: the fold change of expressions) in the transcript abundance and FDR < 0.05 [82]. Identification of unique or overlapping genes within the DEG datasets and the generation of Venn diagrams were determined using Venny 2.1 (https://bioinfogp.cnb.csic.es/tools/venny/index.html).

### GO and KEGG analysis of differentially expressed genes

By searching the NCBI, Uniprot, GO, and KEGG databases, the BLAST (Basic Local Alignment Search Tool) alignment was performed to determine the functional annotation of DEGs. The best matches were selected to annotate the DEGs. GO functional enrichment analysis and KEGG pathway enrichment analysis were carried out by Goatools v0.9.9 [83] and KOBAS v3.0 (http://bioinfo.org/kobas). GO terms and pathway terms with p-value (or FDR) less than 0.05 were considered significantly enriched by differential expressed genes.

## Author Contributions

### Conceptualization

Zhitian Zheng, Huaqi Liu.

### Data curation

Zhitian Zheng, Huaqi Liu, Xiao Luo, Runze Liu, Alexxander Joe, Haolin Li.

### Formal analysis

Zhitian Zheng, Huaqi Liu, Xiao Luo, Runze Liu, Alexxander Joe, Haolin Li, Yanling, Lin.

### Funding acquisition

Zhitian Zheng.

### Investigation

Zhitian Zheng, Huaqi Liu, Xiao Luo, Runze Liu, Alexxander Joe, Haolin Li, Yanling, Lin.

### Methodology

Zhitian Zheng, Huaqi Liu, Haiyan Sun, Yanling, Lin, Yanzhong Li, Yunpeng Wang.

### Project administration

Zhitian Zheng.

### Resources

Zhitian Zheng.

### Software

Zhitian Zheng, Huaqi Liu, Xiao Luo, Runze Liu, Alexxander Joe, Haiyan Sun, Yanling, Lin.

### Supervision

Zhitian Zheng.

### Validation

Zhitian Zheng, Huaqi Liu.

### Writing - original draft

Zhitian Zheng, Huaqi Liu.

### Writing -review & editing

Zhitian Zheng, Haiyan Sun, Yanling, Lin, Yanzhong Li, Yunpeng Wang.

## Acknowledgments

We are grateful for the technical support for Illumina sequencing from the CapitalBio Technology Co., Ltd at Bejing, China and initial data analysis that we received from Biozeron Biotechnology Co., Ltd at Shanghai, China.

## Data availablity

Strains and point mutants of *Fusarium asiaticum* used in this study are available upon request. The raw sequence data from the 36 samples reported in this paper have been deposited in the Genome Sequence Archive (Genomics, Proteomics & Bioinformatics 2024) in National Genomics Data Center (Nucleic Acids Res 2024), China National Center for Bioinformation / Beijing Institute of Genomics, Chinese Academy of Sciences (GSA: CRA014661) that are publicly accessible at https://ngdc.cncb.ac.cn/gsa.

---

**S1 Fig. GO enrichment analysis of DEGs in *F. asiaticum* after 1 μg·mL^-^**^1^ **phenamacril treatment**. The enriched GO categories were shown in a scatter diagram in the comparison groups (A) H1S_1 vs H1S_CK; (B) HA_1 vs HA_CK.(C)HC_1 vs HC_CK; (D) H1R_1 vs H1R_CK. The abscissa represents the enrichment factor, which is the ratio of the number of DEGs enriched in a certain GO term to the number of background genes obtained through sequencing; The ordinate represents the functions enriched by the GO term. The size of small dots represents the number of genes annotated to GO term, the color from red to purple represents the significance of enrichment, and p-value < 0.05 is used as the threshold of significance enrichment for GO enrichment analysis.

**S2 Fig. GO enrichment analysis of DEGs in *F. asiaticum* after 10 μg·mL^-^**^1^ **phenamacril treatment.** The enriched GO categories were shown in a scatter diagram in the comparison groups (A) H1S_10 vs H1S_CK; (B) HA_10 vs HA_CK.(C)HC_10 vs HC_CK; (D) H1R_10 vs H1R_CK.. The abscissa represents the enrichment factor, which is the ratio of the number of DEGs enriched in a certain GO term to the number of background genes obtained through sequencing; The ordinate represents the functions enriched by the GO term. The size of small dots represents the number of genes annotated to GO term, the color from red to purple represents the significance of enrichment, and p-value < 0.05 is used as the threshold of significance enrichment for GO enrichment analysis.

**S3 Fig. KEGG enrichment analysis of DEGs between phenamacril-resistant and -sensitive strains comparison groups.** The top 30 most enriched KEGG pathways were shown in a scatter diagram in the comparison groups (A) HA_CK vs H1S_CK; (B) HC_CK vs H1S_CK; (C) H1R_CK vs H1S_CK. The abscissa represents the ratio of the number of differential genes annotated on KEGG pathways to the total number of differential genes and the ordinate represents KEGG pathways. The size of small dots represents the number of genes annotated to KEGG pathways, the color from red to purple represents the significance of enrichment, and p-value < 0.05 is used as the threshold of significance enrichment for KEGG enrichment analysis.

**S4 Fig. KEGG enrichment analysis of DEGs in *F. asiaticum* after 1 μg·mL^-^**^1^ **phenamacril treatment**. The top 30 most enriched KEGG pathways were shown in a scatter diagram in the comparison groups (A) H1S_1 vs H1S_CK; (B) HA_1 vs HA_CK.(C)HC_1 vs HC_CK; (D) H1R_1 vs H1R_CK. The abscissa represents the ratio of the number of differential genes annotated on KEGG pathways to the total number of differential genes and the ordinate represents KEGG pathways. The size of small dots represents the number of genes annotated to KEGG pathways, the color from red to purple represents the significance of enrichment, and p-value < 0.05 is used as the threshold of significance enrichment for KEGG enrichment analysis.

**S5 Fig. KEGG enrichment analysis of DEGs in *F. asiaticum* after 10 μg·mL^-^**^1^ **phenamacril treatment.** The top 30 most enriched KEGG pathways were shown in a scatter diagram in the comparison groups (A) H1S_10 vs H1S_CK; (B) HA_10 vs HA_CK.(C)HC_10 vs HC_CK; (D) H1R_10 vs H1R_CK.. The abscissa represents the ratio of the number of differential genes annotated on KEGG pathways to the total number of differential genes and the ordinate represents KEGG pathways. The size of small dots represents the number of genes annotated to KEGG pathways, the color from red to purple represents the significance of enrichment, and p-value < 0.05 is used as the threshold of significance enrichment for KEGG enrichment analysis.

**S1 Table. *Fusarium asiaticum* strains and mutants used in this study.**

(DOC)

**S2 Table. A list of primers used in this study.**

(DOC)

